# IRF4 deficiency vulnerates B cell progeny for leukemogenesis via somatically acquired *Jak3* mutations conferring IL-7 hypersensitivity

**DOI:** 10.1101/2022.02.16.480573

**Authors:** Dennis Das Gupta, Christoph Paul, Nadine Samel, Maria Bieringer, Daniel Staudenraus, Federico Marini, Hartmann Raifer, Lisa Menke, Lea Hansal, Bärbel Camara, Edith Roth, Patrick Daum, Michael Wanzel, Marco Mernberger, Andrea Nist, Uta-Maria Bauer, Frederik Helmprobst, Malte Buchholz, Katrin Roth, Lorenz Bastian, Alina M Hartmann, Claudia Baldus, Koichi Ikuta, Andreas Neubauer, Andreas Burchert, Hans-Martin Jäck, Matthias Klein, Tobias Bopp, Thorsten Stiewe, Axel Pagenstecher, Michael Lohoff

## Abstract

The processes leading from disturbed B cell development to adult B cell progenitor acute lymphoblastic leukemia (BCP-ALL) are poorly understood. Here, we describe *Irf4^−/−^* mice as prone to developing BCP-ALL with age. *Irf4^−/−^* preB-I cells exhibited impaired differentiation but enhanced proliferation in response to IL-7, along with reduced retention in the IL-7 providing bone marrow niche due to decreased CXCL12 responsiveness. Thus selected, preB-I cells acquired *Jak3* mutations, probably following irregular AID activity, resulting in malignant transformation. We demonstrate heightened IL-7 sensitivity due to *Jak3* mutants, devise a model to explain it and describe structural and functional similarities to *Jak2* mutations often occurring in human Ph-like ALL. Finally, targeting JAK signaling with Ruxolitinib *in vivo* prolonged survival of mice bearing established *Irf4^−/−^* leukemia. Intriguingly, organ infiltration including leukemic meningeosis was selectively reduced without affecting blood blast counts. In this work, we present spontaneous leukemogenesis following IRF4 deficiency with potential implications for high-risk BCP-ALL in adult humans.

## Introduction

Two signaling pathways via the Interleukin-7 receptor (IL-7R) and the preB cell receptor (preBCR) ensure an orderly progression of B lymphopoiesis.^1–3^ ProB cells adhere to bone marrow (BM) stromal cells (SCs) expressing CXCL12 and VCAM-1 through CXCR4 and VLA-4, respectively,^4^ while SC-derived IL-7 induces their proliferation.^4^ The formation of the preBCR composed of Igµ protein and the surrogate light chain (ψL), consisting of λ5 and VPREB, marks the entrance to the preB cell stage. Signaling via the preBCR in turn induces the transcription factor (TF) interferon regulatory factor 4 (IRF4) which is also critical during T cell differentiation.^5, 6^ In preB cells, IRF4 halts cycling and facilitates recombination of the light chain locus by RAG1/2.^1^ Despite its importance, *Irf4^−/−^* mice still develop, albeit less, surface (s)Igμ^+^ mature B cells,^7^ likely due to a partially redundant function of IRF8. Accordingly, *Irf4,8^−/−^* B progenitors are completely arrested at the preB cell stage.^8^

Disruption of this developmental track can provoke B cell progenitor acute lymphoblastic leukemia (BCP-ALL). In humans, this disease preferentially affects children (age 0-19), while most deaths however occur in the adult population.^9^ Cases affecting adolescents and young adults (AYA) display a different set of driver mutations compared to childhood BCP-ALL.^11–14^

Herein, we report that adult *Irf4^−/−^* mice spontaneously develop BCP-ALL with similarities to human AYA-BCP-ALL and delineate the steps from disturbed *Irf4^−/−^* B lymphopoiesis to overt leukemia.

## Results

### *Irf4^−/−^* mice spontaneously develop preB cell leukemia

Following the serendipitous finding, that some aged *Irf4^−/−^* mice developed tumours and died, we systematically observed 80 *Irf4^−/−^* mice over time. We detected 14 tumours (incidence 17.5 %), that spontaneously appeared in lymph node (LN) areas (mean age: 268d, median: 238d, Fig.1a). Tumours were neither detected in mice younger than 150d nor in C57BL/6 wild-type (wt) mice housed in the same room.

**Fig.1:**
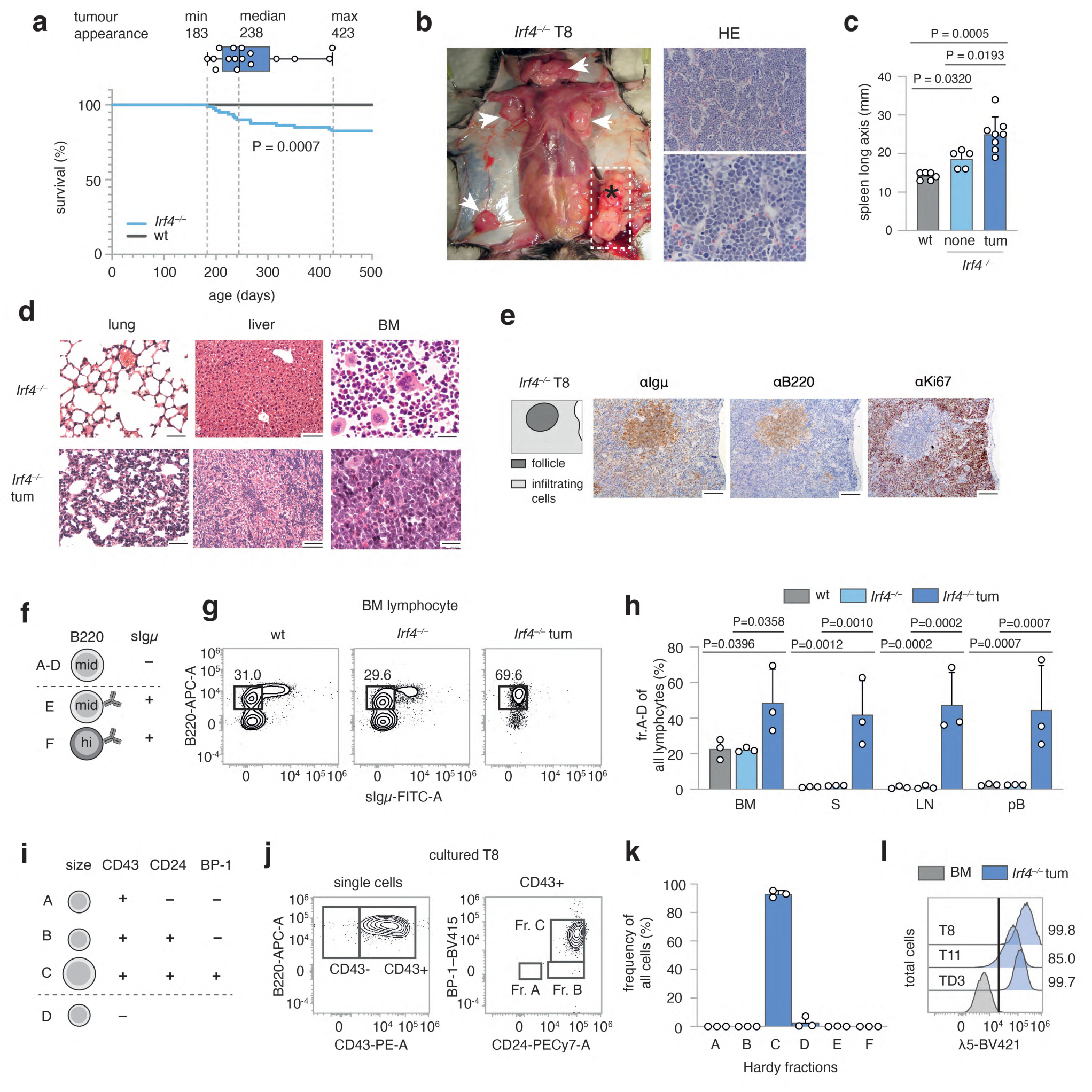
Spontaneous emergence of preB-I cell BCP-ALL in adult *Irf4^−/−^* mice. **a** A cohort of 80 *Irf4*^−/−^ and wt mice was observed over 500 days for tumour development. Kaplan-Meier plot of survival. Box and whisker plot indicates minimum, maximum and median age at tumour appearance of the 14 affected mice. **b** Macroscopic appearance of an exemplary tumour (asterisk) and LNs (arrow heads) in an *Irf4^−/−^* mouse. Right: Hematoxylin-Eosin (HE) staining from the tumour. Scale bars: top: 50 µm, bottom: 20 µm **c** Longitudinal spleen (S) axis (mm) of *Irf4^−/−^* mice with (n = 8) or without (n = 5) tumour and control wt mice (n = 6). tum = tumour **d** HE stainings of lung, liver and BM of tumour mouse T8 and a healthy *Irf4^−/−^* mouse. Scale Bars: 50 µm (lung and liver), 20 µm (BM) **e** IHC-stainings of T8 mouse spleen for Igμ, B220 and Ki67. Scale bars: 100 µm **f** schematic representation of gross Hardy fractioning by surface B220 and Igμ expression **g** whole BM cells from wt, *Irf4^−/−^* and tumour mice were stained for B220 and sIgμ expression and analyzed by flow cytometry **h** quantification of cell frequencies gated as in (**g**) for BM, S, LN and pB (peripheral blood) of n = 3 mice per group **i** tabular representation of Hardy fr.A-D by size, CD43-, CD24- and BP-1- surface expression **j** surface expression of markers as in (**i**) of *in vitro* cultured T8 tumour cells **k** quantification of cell frequencies gated as in (**j**) for three tumours (T8, T11, TD3) **l** surface λ5 expression on T8, T11 and TD3 by flow cytometry in comparison to whole BM cells from *Irf4^−/−^* mice (BM) as negative control. Statistical significance testing was performed with (**c**) one-way Welch-ANOVA followed by Dunnet’s T3 multiple comparison test and (**h**) with two-way ANOVA followed by pair-wise Tukey corrected comparisons within each organ. Bars depict mean ± SD, dots indicate mice (**h**) or distinct *Irf4^−/−^* tumours (**k**)

All tumours (Fig.1b shows a representative tumour *in situ*) were accompanied by lymphadenopathy (arrow heads) and increased spleen size (Fig.1c). Suspected lymphomatous origin was corroborated microscopically (Fig.1b, right panels), with infiltration of mononucleated cells into the BM, lung, and liver (Fig.1d). Due to the known impaired maturation of *Irf4^−/−^* preB cells,^7^ spontaneous eruption of preB-leukemia seemed plausible: In spleen sections (Fig.1e), infiltrating cells stained positive for both B220 and Igμ (although less than untransformed “follicle” B cells) and Ki67. By flow cytometry, BM samples from tumour mice harboured an expanded pro/preB cell compartment (Hardy fraction (fr.)A-D,^21^ B220^mid^sIgμ^−^) (Fig.1f-h). Fr.A-D cells were detected also in peripheral lymphoid organs and blood of tumour mice (Fig.1h). Following the Hardy classification (Fig.1i), we determined tumour cells to be fr.C preB cells (B220^mid^sIgμ^−^CD43^+^CD24^+^BP-1^+^) (Fig.1j-k, sFig.1a). In addition, Igµ, but not Igκ/λ was detected intracellularly (sFig.1b). Lastly, tumours stained positive for surface λ5, part of the ψL (Fig.1l). These attributes characterize the disease as preB-I cell BCP-ALL.

To prove clonality, we sequenced the VDJ junctions of the IgH region in three tumours (supplementary Table 1). Almost all sequences per tumour were identical, demonstrating clonality. The tumours (three examples) further displayed copy number variations (CNV) (sFig.1h), targeting differing genomic regions. Finally, only 500 transferred tumour cells elicited leukemia in wt mice (sFig.1d-e), indicating *bona fide* malignancy.

### B lymphopoiesis in *Irf4^−/−^* mice harbours a hyperproliferative preB-I cell compartment

The uniform appearance of BCP-ALL in *Irf4^−/−^* mice suggested a defined preleukemic pro/preB cell state vulnerable to immortalization. Dimensional reduction of BM samples stained for B cell differentiation markers, identified an enlarged fr.C preB cell compartment already in healthy *Irf4^−/−^* mice (Fig.2a-c). This disturbed, but productive B cell maturation confirms and extends previous reports.^7^ Expression analysis of IL-7Rα and of CD2 (sFig.2b-c), which accompanies cytosolic Igμ expression^22^ further showed an increased frequency of CD2^-/dim^IL-7Rα^+^B220^+^sIgμ^-^ preB cells in *Irf4^−/−^* mice.

**Fig.2:**
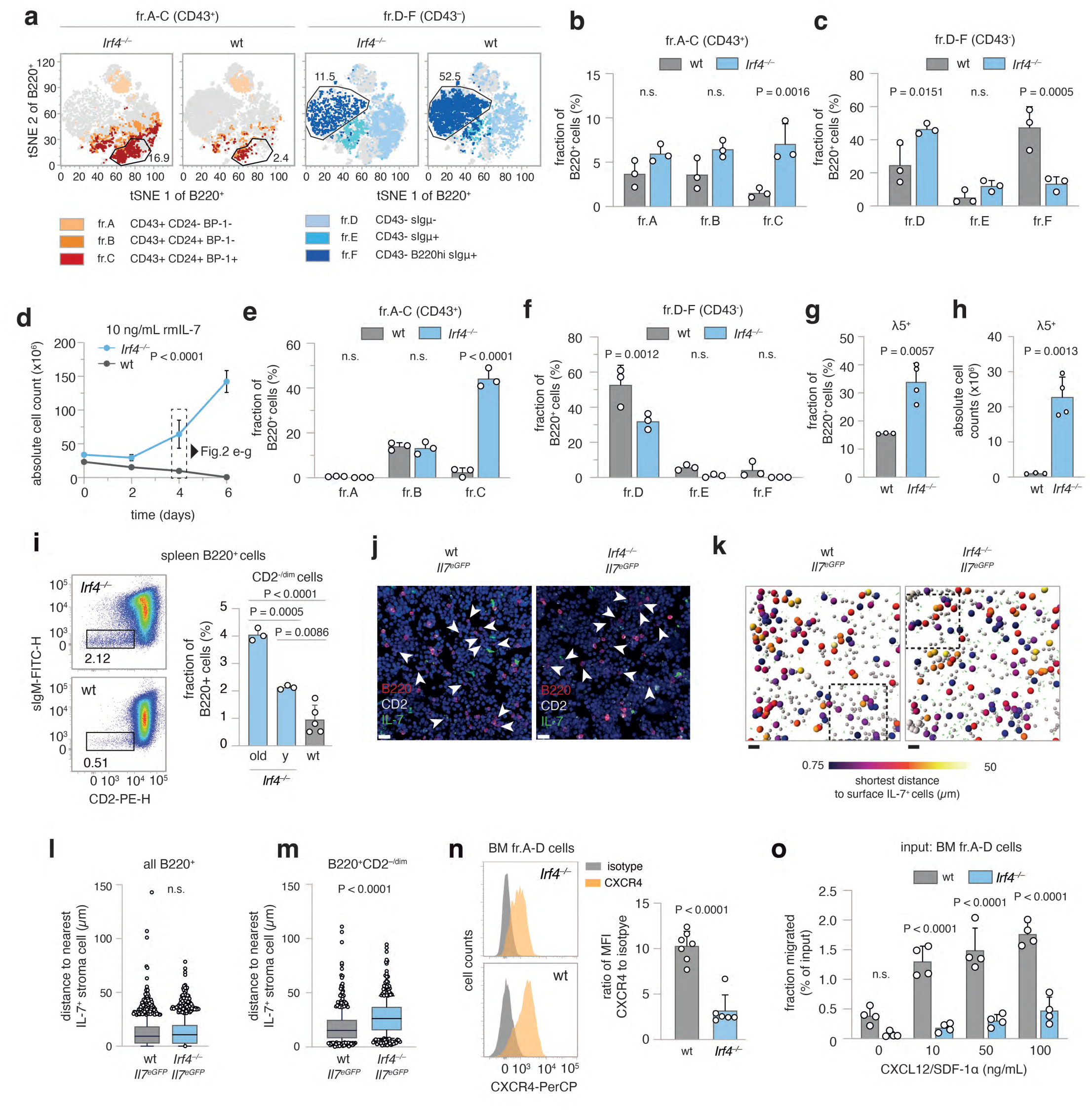
Irf4–/– B lymphopoiesis is preleukemically altered. **a-c** flow cytometric analysis of BM cells for Hardy markers as in Fig.1j. **a** tSNE of BM cells gated on B220^+^ cells. Colours correspond to Hardy fractions identified by the markers detailed in the legend. **b** and **c** quantification of Hardy fraction frequencies for n = 3 mice per genotype. **d-h** BM cells from *Irf4^−/−^* and wt mice were cultured in the presence of 10 ng/mL rmIL-7 for 6 days and **d** counted every two days. **e-f** After 4 days, cells were stained as in (**a-c**) and Hardy fractions quantified. **g** frequency and **h** absolute cell counts of λ5^+^ cells on day 4. **i** spleen cells from *Irf4^−/−^* and wt mice were analyzed for presence of CD2^−/dim^sIgμ^−^ cells within the B220^+^ gate. (y = young) one-way ANOVA, Tukey post-hoc **j-m** 7 µm cryosections from *Irf4^−/−^Il7^eGFP^* and wt *Il7^eGFP^* mice were stained for B220, CD2, GFP and DAPI. **j** exemplary regions of BM cryosections. Arrowheads indicate B220^+^CD2^−/dim^ cells. Scale bars = 15 µm. **k** automated B220^+^ cell detection: grey spheres indicate B220^+^ cells, larger spheres B220^+^CD2^−/dim^ cells, colour-coded for their distance to GFP^+^ cells. Rectangles indicate magnified areas in (**j**). Scale bars = 40 µm. **l-m** quantification of distances to IL-7^+^ cells for **l** all B220^+^ and **m** B220^+^CD2^−/dim^ cells. (n = 4 mice per genotype, one cryosection from femur metaphysis per mouse analyzed). Box and whiskers indicate mean and 95-IQR, dots indicate cells outside 95-IQR. **n** BM cells from *Irf4^−/−^* and wt mice were gated on B220^+^sIgμ^−^ fr.A-D cells and analyzed for CXCR4 expression (left panels as representative staining). Data is presented for n = 7 (wt) and n = 6 (*Irf4^−/−^*) mice as the ratio of geometric mean for CXCR4 to isotype staining.) **o** MACS-purified fr.A-D cells from BM were placed in the top insert of a Boyden chamber and left to migrate towards differing concentrations of CXCL12 for 16 h. Dots represent n = 4 biologically independent experiments, presented as migrated percentage of input cells. Two-Way Anova, Sidak post-hoc for (**b-c**, **e-f, o**), Two-tailed unpaired t-test for (**g-h**, **l-n**)

Purified BM B220^+^ cells from *Irf4^−/−^* and wt mice were cultured with IL-7 (Fig.2d-g) to compare proliferative capacities. After 6d, *Irf4^−/−^* cells had expanded roughly three-fold, whereas wt cell numbers decreased. Phenotypically, *Irf4^−/−^* cells expressed λ5 and accumulated at the fr.C stage (Fig.2e-f); exactly like *Irf4^−/−^* leukemia (Fig.2g). In contrast, wt cells differentiated further, losing surface CD43 (fr.D) (Fig.2e-f) with some cells expressing sIgµ (fr.E). Thus, IL-7 unmasked the leukemic potential of the fr.C compartment in *Irf4^−/−^* mice with both unchecked proliferation and a reinforced differentiation block. Notably, IL-7 dependent *Irf4^−/−^* preB-I cell proliferation was blocked by NIBR3049 and Ruxolitinib, inhibitors of the IL-7R downstream actors JAK3 and JAK1 respectively (sFig.2d).

### *Irf4^−/−^* B cell progenitor exhibit reduced retention to the BM niche

As overt leukemia is characterized by systemic presence, we tested whether already preleukemic *Irf4^−/−^* B cell progenitors would leak from the BM. To reduce the complex Hardy classification, we identified early B cell progenitors, approximately until the preB-I stage, by B220^+^CD2^−/dim^ expression (sFig.2b). Accordingly, we detected higher frequencies of splenic B220^+^CD2^−/dim^ cells in *Irf4^−/−^* than in wt mice (Fig.2h), which accumulated with age. Thus, premature BM evasion adds to the impaired differentiation and hyperproliferation that characterize *Irf4^−/−^* preleukemia.

Potentially, this finding represented a systemic consequence of reduced vicinity to BM niche cells. We therefore analyzed proximity of *Irf4^−/−^* and wt B220^+^CD2^−/dim^ cells to IL-7^+^ BMSCs *in situ* using *Il7*^eGFP^ mice (Fig.2j-m, sFig.2e-f, supplementary movie 1).^23^ In femur cryosections, the B220^+^CD2^−/dim^ subset (Fig.2j, arrowheads) but not the whole *Irf4^−/−^* B220^+^ cell compartment was on average located further away from IL-7^+^ BMSCs, compared to wt control (Fig.2l-m). We excluded differences in IL-7^+^ BMSC abundance between genotypes (sFig.2g).

B progenitor retention to BM is secured via interaction of CXCR4 on pro/preB cells with the IL- 7^+^ BMSC-derived chemokine CXCL12.^24, 25^ Notably, *Irf4^−/−^* pro/preB cells expressed markedly lower levels of CXCR4 compared to wt cells (Fig.2n). Chemokine migration assays with Hardy fr.A-D cells showed that *Irf4^−/−^* cells indeed migrated significantly less towards CXCL12 (Fig.2o). Thus, reduced CXCR4-CXCL12 interaction likely induces the systemic seeding of *Irf4^−/−^* B progeny. Inversely, direct cell interactions are likely not responsible, because *Irf4^−/−^* and wt fr.A-D cells adhered equally to monolayers of OP-9 cells *in vitro* (sFig.2h).

### The IL-7-JAK-STAT-axis is recurrently altered in *Irf4^−/−^* leukemia

Most likely, a second, acquired genetic alteration was necessary for *bona fide* leukemia development and arose with low frequency per time, explaining the affected age and relatively low penetrance. Importantly, IL-7 deprivation of BM-evaded pro/preB cells should create a strong survival stress and potential selection pressure for *bona fide* leukemogenesis. To identify somatically acquired mutations, we performed whole exome sequencing (WES) of three independent tumours (T8, T10, T11) compared to sorted B220^+^mIgM^-^ cells from *Irf4^−/−^* BM. Comparisons of the single nucleotide variants (SNVs) between the three samples identified nine genes affected in all three tumour samples (Fig.3a). Out of these, SNVs in four genes (*Rrs1, Jak3, AW82073* and *Duxf3*) showed alternate base frequencies close to 0.5 or 1 (Fig.3b), suggesting that they could be present on one or both alleles of all leukemic cells. Although we did not exclude oncogenic potential of the other three genes, we focused on *Jak3,* because it is associated with IL-7R signaling. We detected *Jak3* mutations also in tumours TD1, TD2 and T14 (Fig.3c, supplementary Table 2). Thus, six out of six tested tumours carried *Jak3* mutations (“JAK3_mut_”). All mutations targeted either the active kinase domain or the pseudokinase domain regulating JAK3 activity. Some of these SNVs have been described before.^26^ Further, using two different classifiers, no gene fusions could be detected (see methods). Analysis of typical BCP-ALL genes^19^ identified some respective mutations at low frequencies, indicative of subclonal events (Fig. 3d). Among these, mutations in *Jak1*, the partner to JAK3 in IL-7R signaling, were detected in both T8 and T11.

**Fig.3:**
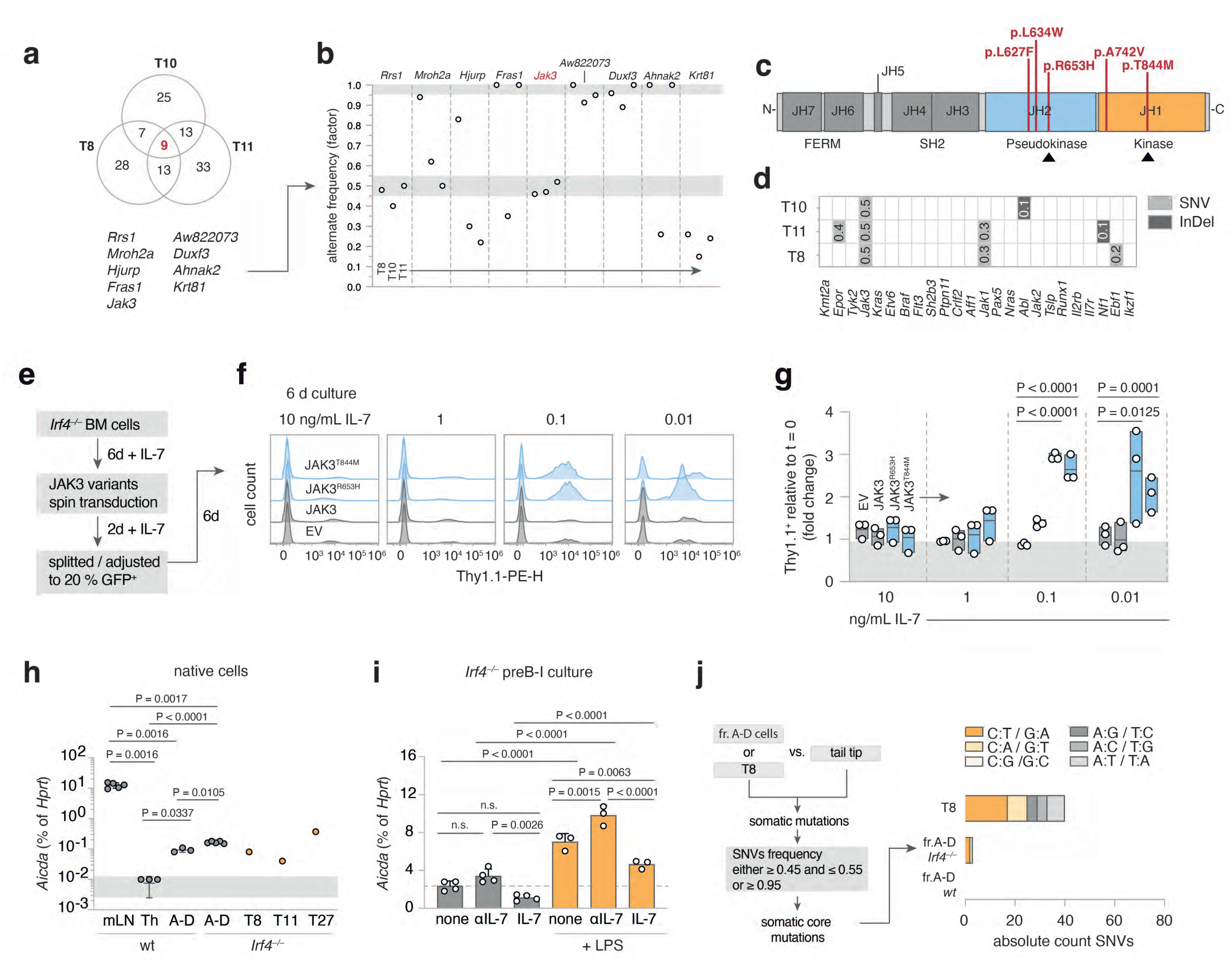
leukemia derived *Jak3* mutations heighten IL-7 sensitivity of *Irf4^−/−^* preB-I cells. **a** Venn diagram of shared mutated genes among WES from three *Irf4^−/−^* leukemia samples (T8, T10, T11) **b** the nine shared genes were filtered for SNV frequency. grey areas: > 0.95 and 0.45-0.55 margins as core mutation filters. **c** the five detected distinct *Jak3* SNVs were mapped onto JAK3 primary structure (JH = Jak homology domain). **d** *Irf4^−/−^* leukemia WES were analyzed for mutations (SNV or InDel = insertions/deletions) in genes commonly altered in human BCP-ALL. Numbers indicate rounded frequencies of alteration. **e-k** *Irf4^−/−^* BM cells were cultured for 6 d with 10 ng/mL rmIL-7, transduced with control or (**e-g**) JAK3_mut_ coding or (**h-k**) STAT5ca coding RVs, rested for 2 days and then split into decreasing IL-7 concentrations. **f** histograms for the Thy1.1 RV infection marker 6 days after splitting. **g** Quantification of Thy1.1^+^ cells after 6 days relative to start of culture (t = 0). Dots indicate n = 3 independent experiments, plotted as floating bars. EV = empty vector. **h** qRT-PCR from wt and *Irf4^−/−^* cells for *Aicda* mRNA expression, relative to *Hprt* expression for n = 3 (Th (= T helper) and sorted fr.A-D wt BM cells), n = 5 (mLN (= mesenteric lymph node) and sorted fr.A-D *Irf4^−/−^* BM cells), n = 1 per tumour T8, T11, T27. **i** *Irf4^−/−^* preB-I cell cultures from whole BM cells were cultured in combinations of IL-7, αIL-7 and LPS for 24 h, as indicated, and analyzed for *Aicda* levels by qRT-PCR for n = 4 (no LPS) and n = 3 (LPS) samples. **j** WES from fr.A-D cells and T8 were compared to tail tip samples to identify SNVs. Filtering on SNV frequency “0.45-0.55 or ≥ 0.95” yielded putative “core mutations”. Absolute numbers of nucleotide exchanges are presented as stacked bars, colours give the type of nucleotide exchange. Two-way ANOVA, Sidak post-hoc for (**h-i**), One-way ANOVA, Tukey post-hoc for (**l-m**)

To analyze the role of the JAK3_mut_, we transduced *Irf4^−/−^* preB-I cell-cultures with retroviruses (RVs) encoding no or wt JAK3 or the JAK3_mut_ R653H and T844M. Culturing transduced cells in the presence of αIL-7 to test for IL-7 independency unexpectedly resulted in cell death after few days with no benefit for cells expressing JAK3_mut_ (sFig.3a-b). To test if JAK3_mut_ would confer advantages with limited IL-7, RV-infected *Irf4^−/−^* preB-I cell-cultures were exposed to decreasing IL-7 concentrations (Fig.3e-g). At 0.1 and 0.01 ng/ml IL-7, both JAK3_mut_-, but not JAK3_wt_-RV led to the outgrowth of transduced over untransduced cells after 6d of culture (Fig.3f-g). Thus, JAK3_mut_ confer IL-7-hypersensitivity, but not -independency. Accordingly, *ex vivo* cultured *Jak3*-mutated T8 and T11 cells also still depended on IL-7 (sFig.3c-d).

### *Aicda* is upregulated in *Irf4^−/−^* preB cells by LPS and deprivation of IL-7

Because five out of six *Jak3* mutations were C to T base exchanges (Table 2), we suspected a specific mutagenic agent. DNA editing enzymes including the APOBEC family member AID^39^ can deamidate cytosines, e.g. during somatic hypermutation.^27, 28^ Repair mechanisms most often ultimately cause C to T conversions.^29, 30^ Notably, AID is induced in wt preB cells by IL-7 withdrawal and LPS stimulation and acts as a facilitator of human BCP-ALL.^31^ Therefore, we compared *Aicda* expression in sorted *Irf4^−/−^* and wt fr.A-D cells to that of wt mesenteric (m)LN- and CD4^+^ T_H_1-cells as controls and to individual leukemia samples. While mLN cells highly expressed *Aicda*, fr.A-D and leukemia-, but not T_H_1-cells, also expressed readily detectable amounts (Fig.3h).

Furthermore, like their wt counterpart,^31^ *in vitro* expanded *Irf4^−/−^* preB-I cells upregulated *Aicda* further under LPS treatment and during IL-7 withdrawal (Fig.3i). This finding can explain how BM evasion and exposure to pathogens might cooperatively initiate mutagenic processes via AID in vulnerable *Irf4^−/−^* preB-I cells.

To test if T8 exhibited signs of previous AID activity on a global level, we analyzed C:T/G:A-transition frequencies in WES of T8, as well as BM-sorted *Irf4^−/−^* and wt fr.A-D cells compared with matched tail-tip samples. Indeed, we found a marked preponderance of C:T/G:A-transitions in T8, when filtering on putative somatic core SNVs (Fig.3j).

### *Jak3* mutations in mice mirror *Jak2* mutations in human Ph-like ALL

Next, we compared *Irf4^−/−^* leukemia to the complex landscape of human BCP-ALL subtypes (reviewed in refs. ^12, 32, 33^), using a published human BCP-ALL cohort for which a random forest classifier had been established (Methods for details).^34^ Only mildly (potentially due to the interspecies comparison) elevated prediction scores were generated for Ph^+^, Ph-like, KMT2a- and DUX4-rearranged human BCP-ALL (sFig.4a-b). Since all of these except Ph-like are defined by specific gene rearrangements, that we had not detected in *Irf4^−/−^* mouse leukemia, we excluded them as comparable candidates.

Ph-like ALL harbours recurrent genetic alterations in signaling molecules, especially in CRLF2 and JAK2.^19^ While BCP-ALL overall preferentially affects children, the incidence of the Ph-like subtype increases from 10 % in children to above 25 % in AYA and adults,^19, 35^ reminiscent of the older age of *Irf4^−/−^* leukemic mice. Furthermore, a published dataset of 154 Ph-like BCP-ALL cases exhibited 10-fold reduced *IRF4* transcripts, when compared to other BCP-ALL subtypes.^19^

While in human Ph-like ALL *Jak2* is commonly mutated, we report recurrent *Jak3* mutations in *Irf4^−/−^* mice. As both proteins are part of distinct but similar signaling complexes in B cell progenitors (Fig.4a), we investigated structural and functional similarities between the specific *Jak3* and *Jak2* mutations. Comparisons of amino-acid-sequences revealed high protein-wide interspecies and intermolecular similarities for both proteins (Fig.4b). Mapping the two amino acids R653 and T844 (mutated in *Irf4^−/−^* mice) onto JAK3 structure predictions, generated by the alpha-fold-algorithm,^36^ revealed that the two amino-acids are in direct contact at an interface of JH1-JH2 domains (Fig.4c). This interface specifically is highly conserved in JAK2 compared to JAK3 (Fig.4d-f, sFig.4c-d). Intriguingly, R683 (corresponding to R653 in JAK3) is by far the most commonly mutated amino-acid in JAK2 in Ph-like ALL, while mutations targeting T875 (corresponding to T844) also have been described.^37^ These findings suggest that mutations in human JAK2 and mouse JAK3 affect a highly similar functional hotspot.

**Fig.4:**
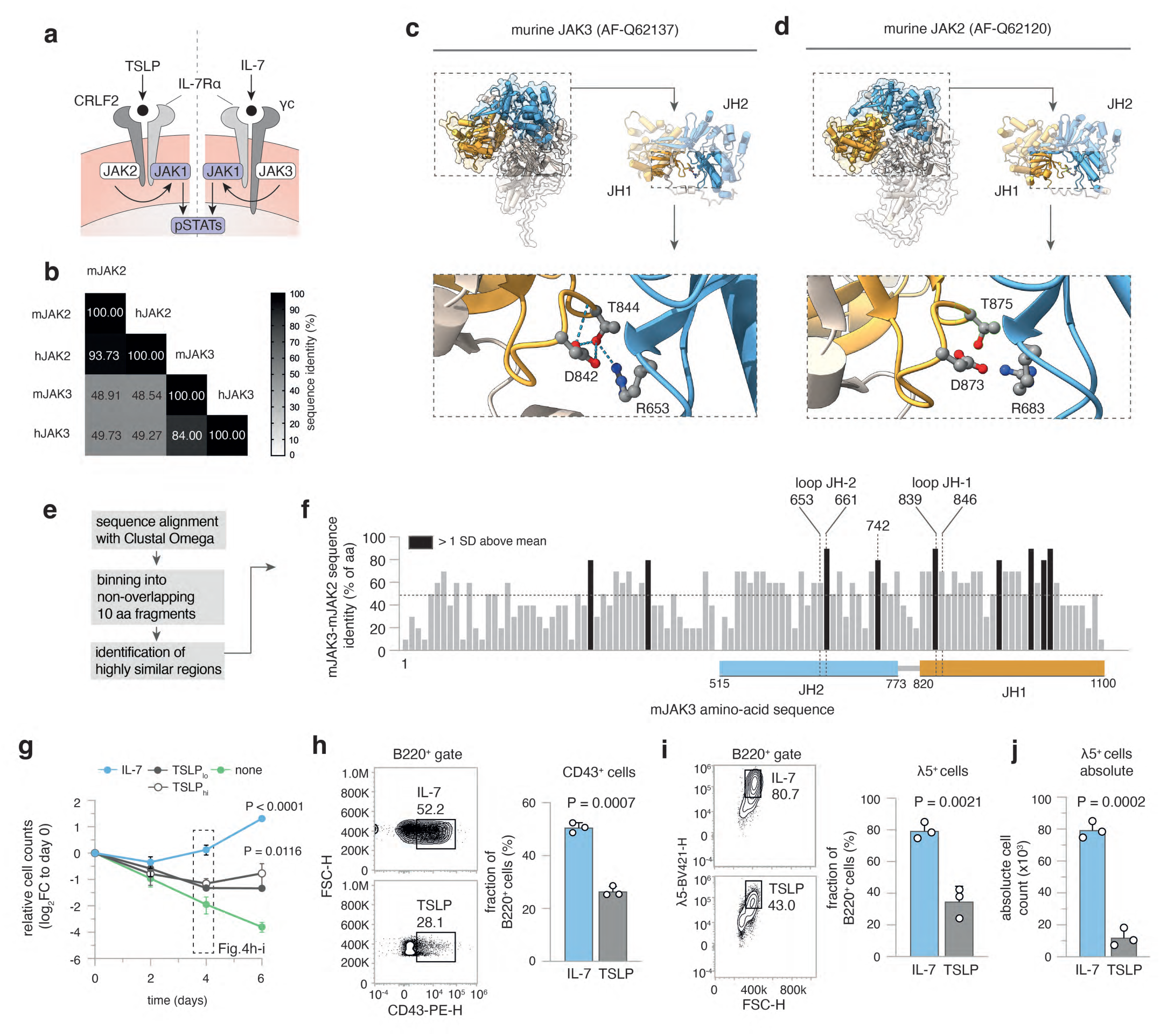
Jak3 mutations heighten IL-7 sensitivity in *Irf4^−/−^* BCP-ALL. **a** cartoon depicting IL-7 and TSLP receptor components. **b** multiple sequence alignment results (using Clustal Omega) of mouse (m) and human (h) JAK3 and JAK2 amino-acid sequences are presented as matrix. Numbers and shade indicate sequence identity as percentage of amino-acids. **c-d** alpha-fold structure predictions of murine **c** JAK3 and **d** JAK2 are presented. Colours indicate domains: orange = JH1, blue = JH2. JH1-JH2 interface is magnified and T844/T875, R653/R683, D842/D873 amino-acids highlighted as ball-and-sticks representations. Dotted lines = hydrogen bonds. **e** overview of analysis for **f**: Sequences of mJAK2 and mJAK3 were aligned using Clustal Omega. The sequence of mJAK3 was binned into 10 non-overlapping amino-acid fragments and the sequence identity to mJAK2 plotted along the mJAK3 sequence. Dotted line = mean protein wide sequence identity, black bars = areas with sequence identity greater than 1 SD above mean. JH1 and JH2 loop regions are mapped onto the sequence, JH1 and JH2 domain regions indicated by coloured rectangles below. **g-j** *Irf4^−/−^* BM cells were cultured for 6 days in the presence of 10 ng/mL rmIL-7, 10 or 100 ng/mL rmTSLP(_lo/hi_) or no cytokine (none). **g** log_2_ of cell counts relative to day 0 for n = 3 independent experiments plotted as means ± SD. One-way ANOVA, Sidak post-hoc comparing cytokine effect **h-i** On day 4, **h** CD43^+^ and **i** λ5^+^ cells within B220^+^ cells were recorded for IL-7 and TSLP treated cultures. Numbers indicate percentages within the depicted gates of B220^+^ cells. **j** absolute counts of λ5^+^ cells at day 4. Dots in (**h-j**) indicate n = 3 independent experiments, presented as bars (mean ± SD). Unpaired two-tailed t-test for (**h-j**)

As mentioned above, JAK2 and JAK3 are part of distinct, but similar receptors: JAK3 binds the common γ-chain involved in IL-7 signaling, while JAK2 associates with CRLF2 involved in TSLP signaling. Both signals involve the IL-7Rα chain and the same downstream pathways (STATs, PI3K).^38^ Therefore, alternative presence of JAK3/JAK2 mutations between mouse and human BCP-ALL might reflect different cytokine preferences. Human proB/preB cells proliferate in response to both TSLP and IL-7.^39^ However, in *Irf4^−/−^* BM cells IL-7, but not TSLP induced robust proliferation (Fig.4g) as well as high frequencies and absolute counts of CD43^+^ (Fig.4h) and λ5^+^ preB cells (Fig.4i-j).

### IRF4 re-expression leads to cell death and differentiation

As *Irf4* deletion was a prerequisite for leukemia in our model, we examined the effect of forced IRF4 re-expression, using RVs coding for GFP alone (EV-RV) or plus IRF4 (IRF4-RV). When re-introducing IRF4 into T8 or T11, GFP^+^ IRF4-expressing-, but not GFP^+^ control-cells gradually disappeared over time (Fig.5a). AnnexinV/PI stainings confirmed apoptosis (not shown). Further, we noted loss of surface λ5-expression induced by IRF4-RV (Fig.5b-c). Comparing the transcriptomes of still viable cells 24h after transduction revealed strong induction of “apoptotic process” and “innate immune response” gene ontology (GO) gene sets (gs) (Fig.5d). Markov clustering of GO gs affected by IRF4 re-expression (Fig.5e-f) further identified several coregulated B cell differentiation gs (Fig.5f), with downregulated ψL components *Igll1*, *Vpreb1* and *Vpreb2*, but upregulated differentiation genes including *Igμ*, *Igκ* and *Blnk* (Fig.5g-h, sFig.5a-b). Similar results were obtained for T11 (sFig.5c-d). Therefore, fully transformed leukemia remained targetable by IRF4 re-expression.

**Fig.5:**
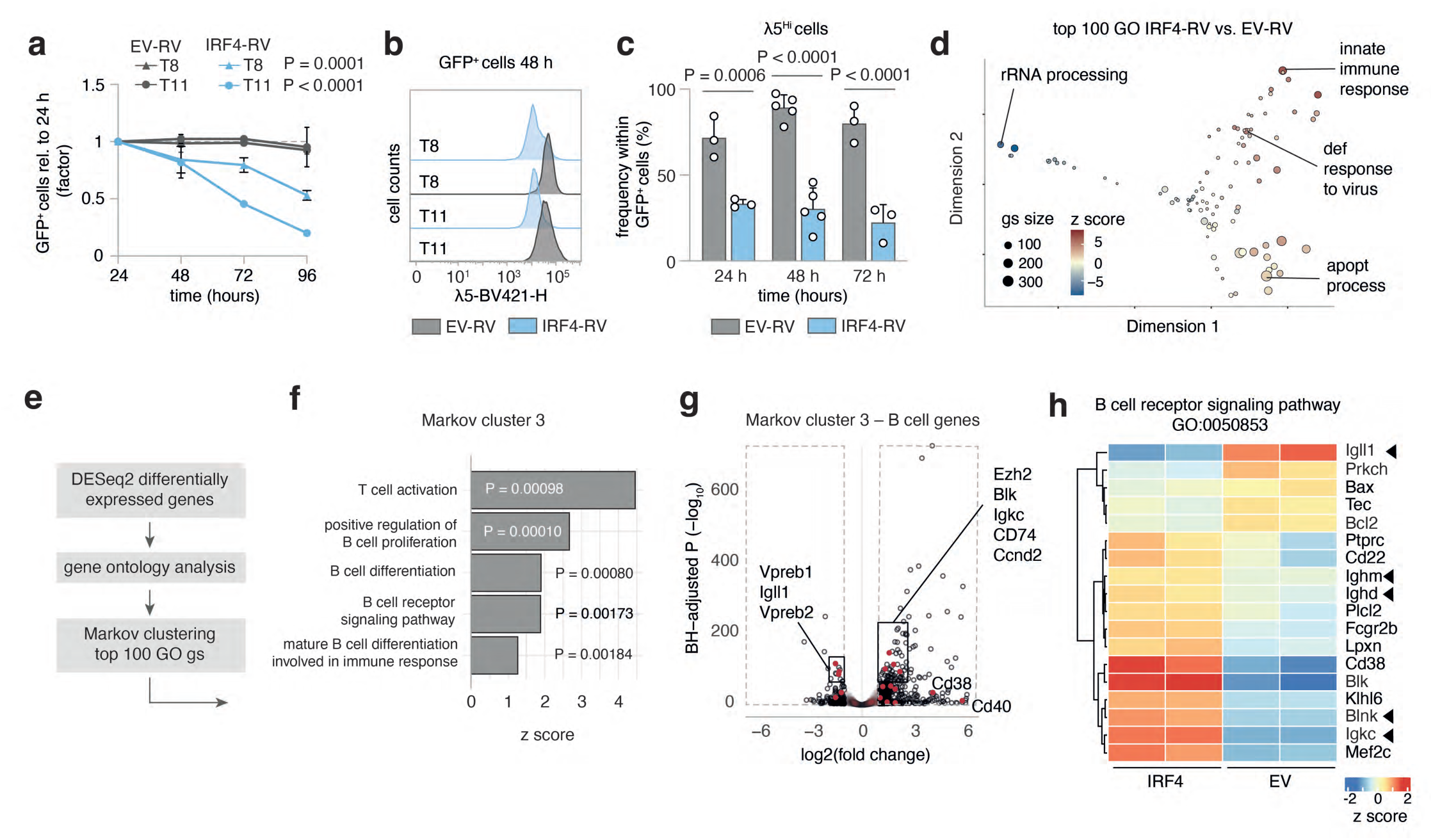
IRF4 re-expression results in apoptosis and differentiation of leukemia cells. **a-c** T8 and T11 cells were transduced with IRF4-RV or control empty vector (EV)-RV. **a** GFP^+^ cell frequency normalized to 24 h after transduction was recorded. One-Way ANOVA, Sidak post-hoc for RV effect per tumour. Mean ± SD of n = 3 independent experiments. **b** representative histogram of λ5 surface expression of GFP+ cells at 48 h. **c** Pooled Quantification of λ5^Hi^ cells for three (24, 72h) to five (48h) independent experiments for T8 and T11. **d-h** T8 cells were collected in duplicates at 24 h after EV-RV and IRF4-RV transduction and subjected to bulk RNAseq. **d** MDS plot of top 100 gene ontology (GO) gene-sets varying between EV and IRF4 transduced T8. Representative gene-sets annotated. Size of circles = number of genes, colour = z score. **e** analysis strategy for GO gene-set clustering using Markov clustering. **f** gene-sets from Markov cluster 3 and corresponding P values and z scores. **g** volcano plot of B cell genes from Markov cluster 3 (red) highlighted within all differentially regulated genes (black). **h** heat map of B cell receptor signaling GO gene-sets. Immunoglobulin genes and the tumour suppressor *Blnk* are marked. Colour = z score.

### Small compound agents affecting *Irf4^−/−^* leukemia *cells in vitro*

Next, we screened a collection of kinase inhibitors for their capacity to kill *Irf4^−/−^* leukemia cells *in vitro*. We included NIBR3049 targeting JAK3, Ruxolitinib, an inhibitor of JAK1/2 (downstream of JAK3) and Dexamethasone, a cornerstone for treating lymphomatous malignancies. Furthermore, we included inhibitors of NFκB (IKK, TAK1), JNK, MEK, ERK, PP2A, GFI1, FAK and of the Bruton tyrosine kinase (BTK) acting downstream of the BCR.

A variety of these substances potently killed tumour cells (sFig.6a), implying involvement of multiple pathways in leukemia cell survival. Efficacy of Ruxolitinib and NIBR3049 corroborated our results concerning *Jak3* driver mutations. Furthermore, inhibitors of GFI1 and PP2A as well as NFκB and JNK were potent. In contrast, inhibiting BTK, MEK and ERK had no impact.^40^

### *In vivo* therapy of established *Irf4^−/−^* B-ALL

Next, we implemented JAK inhibition as *in vivo* treatment for *Irf4^−/−^* leukemia. We began induction therapy with Dexamethasone around day 12 after adoptively transferring 3x10^5^ T8.1 cells *i.p*. into wt mice (Fig.6a), when overt leukemia was noted in peripheral blood (pB) (Fig.6b “pre”). After 7d of treatment, leukemic cell numbers in pB were robustly reduced (Fig.6b “post”), although few cells reproducibly remained detectable (Fig.6c). Maintenance therapy was continued with Ruxolitinib or vehicle control by oral gavage twice daily for the following 12 days (Fig.6a,c). Importantly, the half-life of Ruxolitinib in mice is only 0.8 h (“Australian Public Assessment Report for Ruxolitinib”, Australian Government), implying that any observed *in vivo* effectiveness might be underestimated.

**Fig.6:**
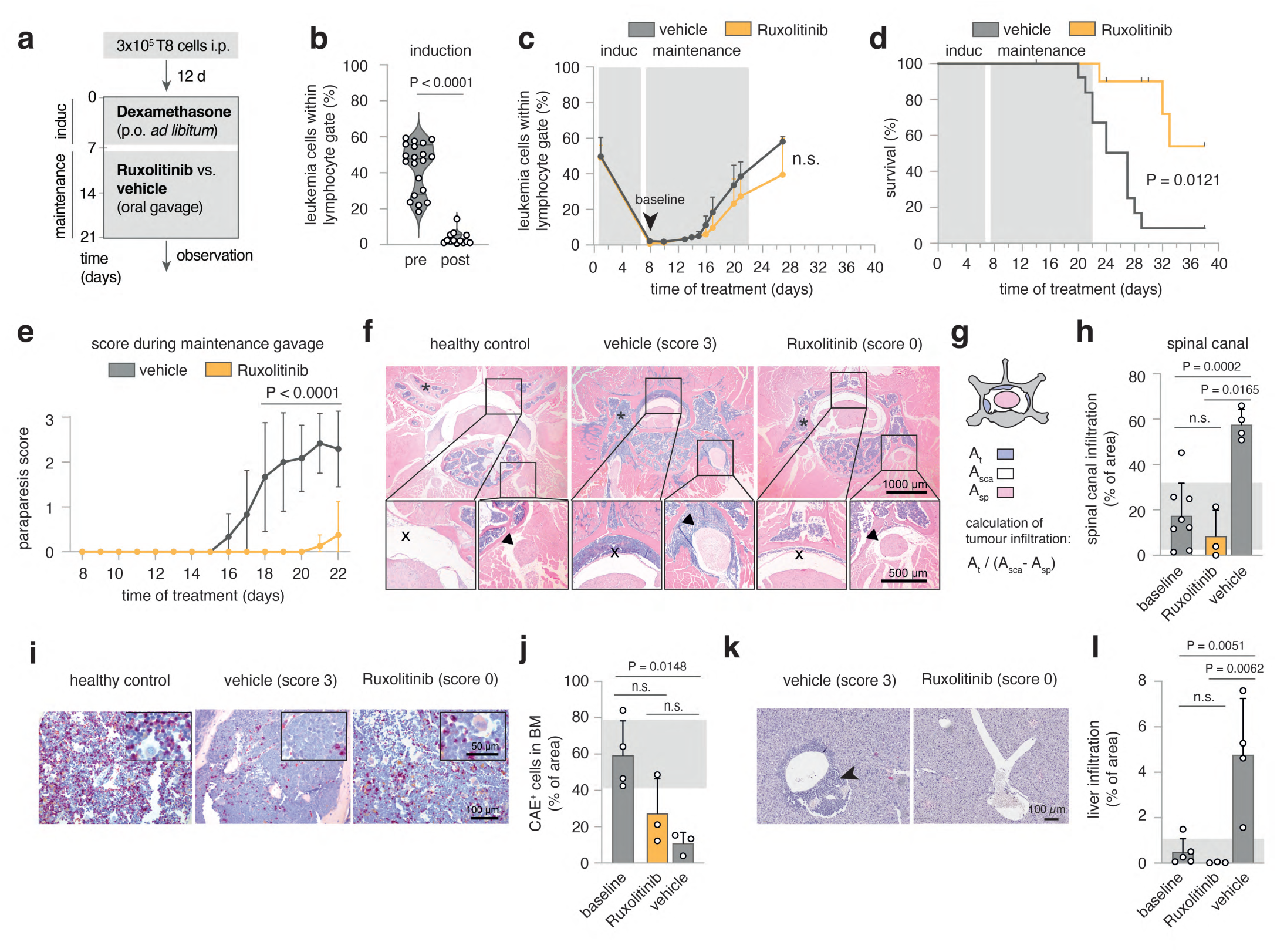
Ruxolitinib reduces leukemic meningeosis and organ infiltration *in vivo*. **a** schematic overview of experimental design. Day 0: injection of mice with 2x10^5^ T8 cells. After 12 days initiation of Dexamethasone induction therapy supplied in drinking water for seven days. Maintenance therapy comprised either Ruxolitinib-phosphate (11 mice) or vehicle control gavage (13 mice) twice daily for 14 days. Mice were scored daily and blood sampling performed regularly. **b** leukemia cell frequency (B220^+^sIgμ^-^) within lymphocyte gate before and after induction with Dexamethasone. Two-tailed unpaired t-test **c** time-course of leukemia cell frequencies in peripheral blood for Ruxolitinib and vehicle treated mice. **d** Survival as Kaplan-Meier plot analyzed with Log-rank test. In the Ruxolitinib group, 4 mice were excluded and censored due to intervention related adverse reactions or due to their use for to the analysis described in (**f-l**). **e** Disease scores, determined as described in methods. Mean ± SD of the scores per indicated treatment group analyzed by two-way ANOVA, Sidak post-hoc. n = 2 replicate experiments for (**b-e**) with similar outcome. **f** Exemplary histopathology (HE) of healthy or leukemia bearing mice (score 3, vehicle-treated or score 0, Ruxolitinib-treated). One representative mouse per condition. Bar size in the bottom right corners. Top panels: overview of cross-sectioned lumbar vertebrum, bottom inserts from spinal canal (left) and spinal nerve root (right). **g** Schematic representation of calculation of tumour infiltration into the spinal canal. (A_t_: area of tumour infiltration, A_sca_: area of total spinal canal, A_sp_: area of the spinal cord) **h** Quantification of spinal canal infiltration according to (**g**) for n = 8 after induction (baseline), n = 3 score 0 (Ruxolitinib) and n = 4 score 3 (vehicle) mice. **i** Representative CAE stainings from vertebral BM for score 0 and score 3 mice. **j** Quantification of area occupied by CAE^+^ cells relative to total BM area for n = 4 (baseline), n = 3 (score 0, Ruxolitinib) and n = 4 (score 3, vehicle) mice. **k** Representative HE stainings from liver tissue for score 0 and score 3 mice, Scale bar bottom right. **l** Quantification of tumour infiltrated area relative to whole liver area for n = 5 (baseline), n = 3 (score 0, Ruxolitinib) and n = 4 (score 3, vehicle) mice. Each dot represents measurements of three complete liver cross-sections per mouse.

Despite maintenance therapy, leukemic cells in pB reappeared, with no significant difference between treatment groups (Fig.6c). However, treatment with Ruxolitinib resulted in a clear survival benefit (Fig.6d) and marked improvement of a prominent neurological symptom: in sham-treated animals, temporary limpness of tail and hind legs occurred seconds after gavage, which we quantified using a newly established scoring system (ranging from 0 to 3, see Methods).

Mechanistically, ultrasound revealed an echogenic paravertebral mass (sFig.7a-b) in score 3, but not score 0 mice. By histology, score 3 correlated with severe infiltration of blasts into the spinal canal (X in Fig.6f), extending into spinal nerve roots (arrowhead in Fig.6f). Therefore, paraparesis likely represented a manifestation of mouse leukemic meningeosis, exacerbated by gavage-induced increases in intraabdominal pressure.

Paraparesis was reproducibly relieved during Ruxolitinib treatment (Fig.6e), correlating with the suppression of perimyelon infiltration that ensued in vehicle-treated mice after the end of induction therapy (Fig.6h). In contrast, the severely impaired hematopoiesis in sham-treated mice, indicated by low CAE^+^ cell frequencies, was not significantly ameliorated by Ruxolitinib (Fig. 6i-j).

These findings raised the possibility that Ruxolitinib preferentially targets infiltration of solid organs rather than BM or pB. Accordingly, Ruxolitinib fully blocked the liver infiltration as observed in sham-treated mice (Fig.6k-l). As tissue infiltration is regulated by homing receptors, we treated T8.1 and T8.2 cells with Ruxolitinib *in vitro* and recorded the expression of CD29 (integrin β1), which pairs with various integrin alpha chains involved in cell- and tissue-adhesion.^41, 42^ Notably, on T8.1 and T8.2, Ruxolitinib reduced CD29 expression dose-dependently (sFig.6c-e) while even slightly increasing expression of MHC I molecules (H2D^b^, H2K^b^), stained as specificity control.

## Discussion

The herein described spontaneous leukemogenesis in *Irf4^−/−^* mouse stresses the particular vulnerability of preB-I cells. Our data provides insights for a.) conditions promoting leukemogenesis, b.) functional consequences of *Jak* mutations, c.) parallels of mouse and human BCP-ALL and d.) potential *in vivo* treatment:

a.) We provide evidence for a two-hit leukemogenesis model: The first hit (*Irf4* loss) resulted in reduced differentiation, IL-7 dependent hyperproliferation and impaired retention to the BM niche (Fig.7). A second hit (targeting *Jak3* in our model) created a dominant survival signal, resulting in overt preB-I leukemia.

**Fig.7:**
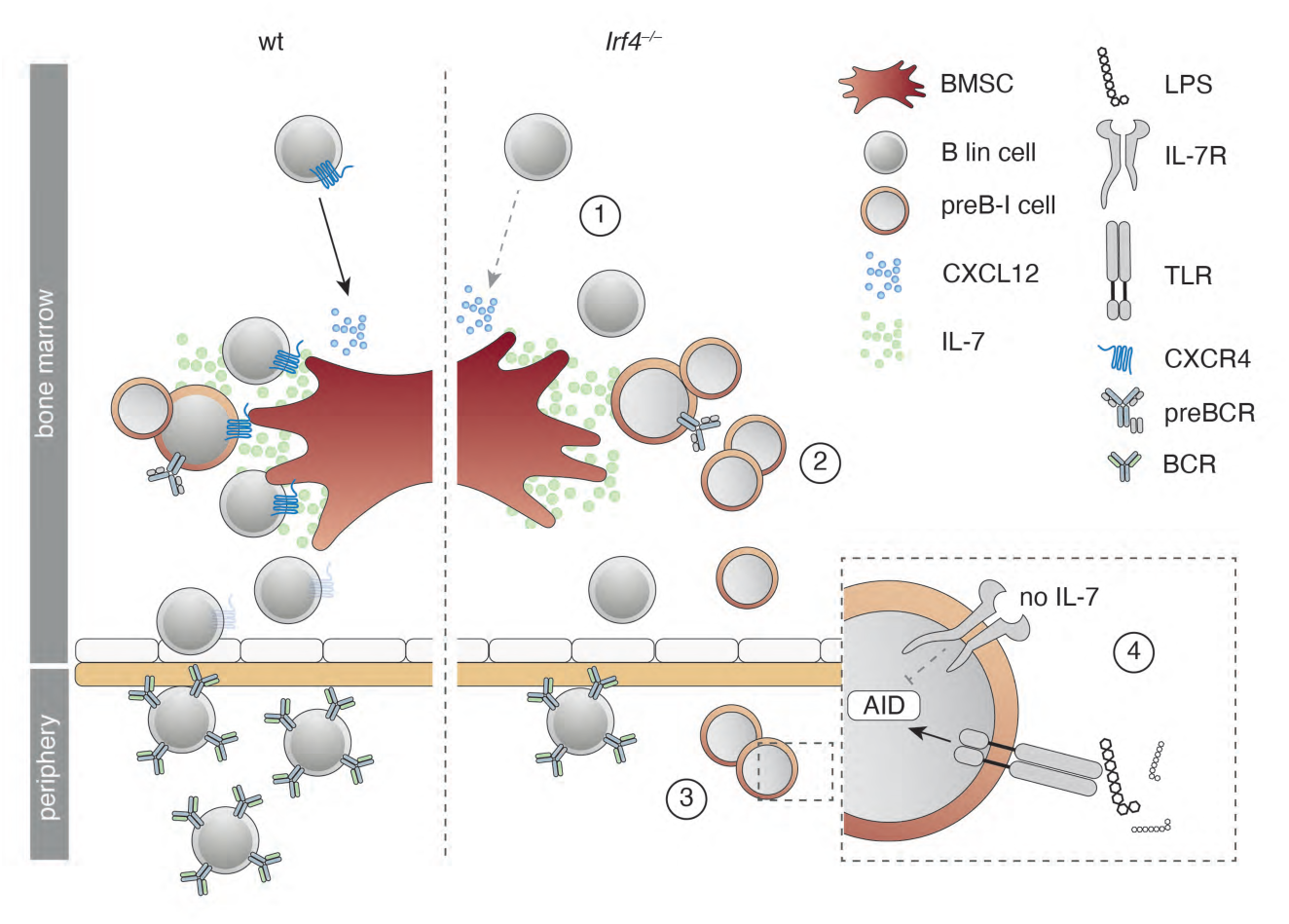
Summary of the preB-I preleukemic state induced by IRF4 deficiency. Cartoon summarizing the findings for IRF4 deficient compared to wt B lymphopoiesis. B lineage (lin) cells are less responsive to BMSC derived CXCL12 due to reduced surface CXCR4 expression (1). *Irf4^−/−^* preB-I cells exhibit impaired differentiation and IL-7 dependent hyperproliferation (2). *Irf4^−/−^* preB cells escape into the periphery (3), where a combination of IL-7 deprivation and danger associated molecular patterns (such as LPS) might induce AID expression (4), fueling mutagenesis.

The induction of BCP-ALL in *Irf4^−/−^* mice is similar to *Ikzf1* and *Pax5* mutated mouse models,^43–, 45^ implying similarities between these TF-alterations. Probably, one shared mechanism is the differentiative impairment. Importantly, for *Irf4^−/−^* fr.A-D cells we even detect slightly higher levels of *Pax5* compared to wt fr.A-D cells (sFig.8), ruling out that the findings in *Irf4^−/−^* mice merely mirror those of *Pax5* deficiency. The reverse remains conceivable; that *Ikzf1* and *Pax5* mutations converge lowering IRF4 expression.

In addition to mice mutated in *Pax5* or *Ikzf1*, *Irf4/Irf8^−/−^* and *Irf4/Spi1^−/−^* mice have been shown to develop leukemia early in life at high incidence.^20, 46^ Contrasting these studies, we report that single deficiency for IRF4 fully suffices for leukemogenesis. We excluded secondary alterations in *Irf8, Spi1* in our model: we found unchanged expression and gene sequence of IRF8 (not shown) and normal amounts of *Spi1* transcripts (sFig.8a) in *Irf4^−/−^* fr.A-D cells. The single IRF4 deficiency models potential clonal initiating events better than *Irf4/Irf8^−/−^* or *Irf4/Spi1^−/−^* mice, because *Irf4^−/−^* mice harbour productive B cell development.

We newly describe that a preleukemic alteration can lead to reduced BM retention, presenting a tentative explanation for the induction of mutagenic signals, as deprivation from IL-7 and exposure to bacterial compounds can cooperatively induce the mutagenic agent AID.^31^

b.) Why do *Jak3* mutations only lead to enhanced sensitivity to, but not complete independence of IL-7? Analysis of JAK3 and JAK2 structure implied that mutations of R683/R653 and T875/T844 might decrease JH1-JH2 interaction strength. This would imply reduced auto-inhibition as the GOF mechanism – in line with findings for the JAK family member TYK2.^48^ This alone cannot explain cytokine independency, owing to the receptor biology: The two preassembled receptor-chains keep JAKs intracellularly separated.^49^ Ligand binding is needed for a conformational change that brings JAKs into the proximity needed for cross-phosphorylation.

To explain our observations, we propose an oncogene model with two equilibria (Fig.8a-b): the first is determined by cytokine concentration and dictates the probability of receptor conformation change (Fig.8a). The second, independent equilibrium (Fig.8b), is determined by the interaction strength at the JH1-JH2 interface and dictates the probability of JH1 and JH2 dissociation. Only the combination of the “bound” and “active” state (Fig.8c) would result in the elicitation of a signal (Fig.8c, green frame).

**Fig.8:**
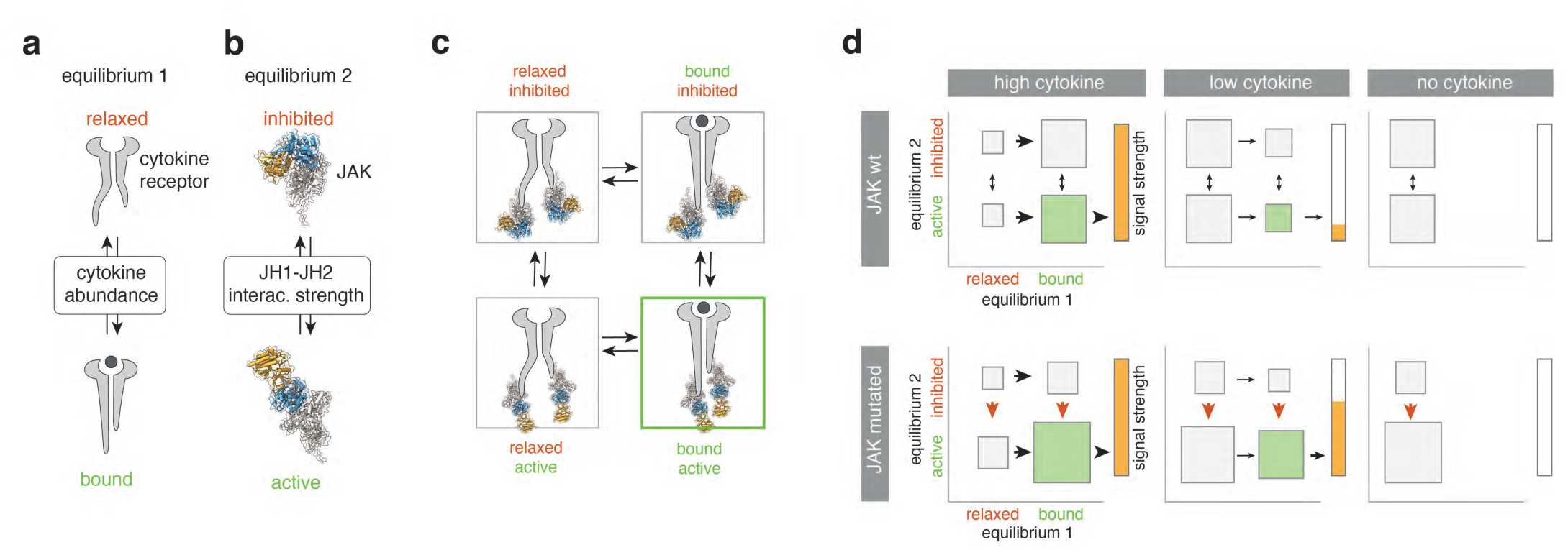
A “two-equilibrium model” explains JAK mutant effects in primary preB-I cells. **a** Equilibrium 1 is determined by cytokine abundance and dictates the cytokine receptor state (bound vs. relaxed). **b** Equilibrium 2 is determined by the interaction strength between JH1 and JH2 domains in JAKs and dictates the JAK state (inhibited vs. active). **c** Equilibria 1 and 2 interact to create four possible states. Green lettering indicates a signaling favoring state. Green frame indicates the actively signaling state. **d** For high (left), low (middle) and no (right) cytokine in the presence or absence of JAK3 mutations, hypothetical probabilities of equilibria states as in (**c**) are presented. Size of rectangles signifies likelihood of state relative to others. Arrows indicate shifts of equilibria, red arrows indicate the effect of JAK mutations. Green rectangle = active signaling state (bound and active). Bars to the right of each panel indicate signal strength as a direct result of the two equilibria adjacent to it.

In this model, JAK mutations would only affect the second equilibrium (Fig.8d, red arrows). Therefore, sporadic ligand binding would still be needed for elicitation of signaling. The model stringently predicts the better exploitation of low cytokine concentrations for JAK_mut_ that we observed *in vitro*. Fig.8d depicts theoretical probabilities of receptor states for varying cytokine concentrations in the presence (top row) or absence (bottom row) of JAK mutations.

Our findings that JAK3_mut_ confer heightened cytokine-sensitivity, but not -independence, is in contrast to what has been found for JAK2-R683G mutants expressed in the commonly used BaF3 cell-line.^51^ However, BaF3 cells depend on IL-3 and not IL-7Rα cytokines (i.e. IL-7 or TSLP). Therefore, it remains conceivable, that the IL-3 receptor provides a different physiology, which may deviate from the IL-7R physiology in primary B progenitors.

c.) The finding that *Irf4^−/−^* preB-I cells respond preferentially to IL-7 over TSLP presents a possible explanation, why mouse models of BCP-ALL acquire *Jak3* mutations, whereas the human Ph-like disease typically harbours *Jak2* mutations. Our comparison of JAK structure predictions yielded corresponding mutations likely to elicit similar downstream effects.

d.) Lastly, our *in vivo* experiments reinforce Ruxolitinib as a potential treatment for JAK driven BCP-ALL. The compound represents an important therapeutic agent in myeloproliferative disease and is already studied for treatment of Ph-like-ALL.^52, 53^ We describe a preferential effect of Ruxolitinib on CNS- and organ infiltration, potentially due to reductions in integrin expression on leukemia cells. These effects are of translational importance because current CNS-targeted therapies for ALL remain toxic. Future studies should therefore evaluate efficacy of Ruxolitinib for leukemic organ infiltration in other leukemia models.

## Methods

### Mice

C57Bl/6 mice were purchased from Charles River, Sulzfeld, Germany. *Irf4^−/−^* mice^7^ and *Il7^eGFP^* mice^55^ (provided by Koji Tokoyoda, DRFZ Berlin) were bred on the C57Bl/6 background and housed in the animal facility of the Biomedical Research Center at the university of Marburg, Germany. If not stated otherwise, all mice used in the presented experiments were 8-12 weeks old and sex-matched.

### Tumor cell lines and cell culture

Stable tumor cell lines T8.1, T8.2 and T11 were established from primary *Irf4^−/−^* leukemia cells (derived from primary tumour 8, i.e. T8, or tumour 11) by culturing them on a monolayer of irradiated (30 Gy) ST2 stromal cells^56^ grown to confluency in Opti-MEM medium (31985070, ThermoFisher Scientific) supplied with 1 % cell culture supernatant from JIL-7.6 J558 cells^57^ (a gift from Fritz Melchers, Berlin) as a source of IL-7. After several passages, T8 and T11 cells grew independently of ST-2 cells. For *in vitro* inhibitor experiments, 2.5 x 10^5^ T8.1 or T8.2 cells (or T11 cells) were cultured in 500 µL RPMI medium in 48 well plates in the presence of the indicated concentrations of inhibitors. To determine the percentage of viable cells, samples were stained using Annexin V and propidium iodide (PI) (see below) after 48 h. Substances used include: Defactinib (S7654, Selleckchem), Oxocaenol (O9890, Sigma), GANT61 (Sigma, G9048), SP203580 (EI-286-0001, Enzo), SP600125 (EI-305-0010, Enzo), PD98059, Promega), Ibrutinib (S2680, Selleckchem), BAY11-7082 (ALX-270-219, Alexis), Dexamethasone (PZN 08704491, mibe GmbH) and Ocadaic acid (O4511, Sigma).

### Murine pro/preB cell cultures

Femur and tibia bones from 8-12 weeks old mice were explanted and cleaned from adherent tissues. Cells were extracted via centrifugation at 11 x 10^3^ RPM for 10 s. Total BM cells were enriched for B220^+^ (sIgµ^-^) B lineage cells using an inhouse magnetic activated cell sorting protocol. Briefly, whole bone marrow cells were stained with a mix of FITC-conjugated antibodies to (Igµ), CD11b, B220, Ter119, CD49b, CD4 and CD8 (all from eBioscience), followed by incubation with an anti FITC/streptavidin/biotin/magnetic bead complex (Miltenyi Biotec) and magnetic sorting using a microcentrifugation tube stand (Miltenyi Biotec).^58^ Sorting efficiency, as confirmed by flow cytometry, routinely exceeded 90 %. Cells were seeded at a density of 1 x 10^5^ cells per well in 200 µL RPMI complete (96-well plates, Greiner). Pro/preB cell cultures were propagated with 10 ng/mL rmIL-7 (217-17, Peprotech) in RPMI-1640 medium complete (R8758, Sigma-Aldrich, supplemented with: 10% FCS (Sigma-Aldrich), 2mM L-glutamine (Biochrom), 50 µM #-mercaptoethanole (Sigma-Aldrich), 0,03/0,05 g per 500 mL Penicillin G/Streptomycin Sulfate, 1 % non-essential amino acids (PAA Laboratories)). In some experiments, pro/preB cells (1.25 x 10^6^/mL medium) were treated for 24 h with LPS (Sigma, 1 µg/ml), anti-IL-7 (BioXCell, 10 µg/ml), rmIL-7 or respective combinations, before generating mRNA for qRT-PCR.

### Transwell migration assay and OP-9 adhesion assay

For transwell migration assay, Hardy fr.A-D cells were magnetically sorted from BM of wt and *Irf4^−/−^* mice as described above (with addition of FITC-conjugated anti-Igµ antibody), and 2 x 10^5^ cells in RPMI (without additives, FCS-free) 10 ng/mL rmIL-7 seeded in 50 µL in the top chamber of 96 well 5 µm pore uncoated 96 well transwell plates (HTS transwell^®^ Corning). The bottom chamber was flooded with 200 µL RPMI containing indicated concentrations of rmCXCL12 (Peprotech). After 16 h, inserts were removed, cells in the bottom chamber collected, counted and analyzed for B220 surface expression using flow cytometry. The fraction of migrated cells was calculated as n(migrated) * freq_B220_(migrated) / n(input) * freq_B220_(input). Normalization to B220^+^ cells reduced interexperimental differences due to differences in cell purity after magnetic selection. For OP-9 adhesion assays, 5 x 10^3^ OP-9 cells (a gift from Hyun-Dong Chang, DRZF Berlin) were seeded in 96 well microtiter plates 24 h before the assay. At the day of the assay, fr.A-D cells were purified as above and 2 x 10^5^ fr.A-D cells seeded on top of OP-9 monolayers in RPMI complete + 10 ng/mL rmIL-7. Plates were centrifuged briefly to accelerate cell descension. After 1 h, suspended cells were collected in the supernatant and by washing OP-9 monolayers two times with PBS.

### Flow cytometry and cell sorting

For surface staining of B lineage markers, cells were harvested, resuspended in PBS/1% FCS and stained with anti-B220 (RA3-6B2, Biolegend), anti-Igµ (II/41, BD Bioscience), anti-CD43 (RM2-5, Biolegend), anti-CD24 (M1/69, invitrogen), anti-BP-1 (BP-1, BD Bioscience), anti-CD2 (RM2-5, Biolegend), anti-CXCR4 (L276F12, Biolegend), anti-CD127 (=IL-7Rα) (A7R34, BD Bioscience), anti-CD179b (=λ5) (LM34, BD Bioscience) as indicated (20 min at room temperature in the dark). All antibodies were employed at a dilution of 1:500. Fluorescence was recorded using either a FACS Aria III (BD) or an Attune NxT (Thermo-Fisher) analyzer. Data analysis was performed using the FlowJo V10 software (BD). For dimensional reduction we used the t-Distributed Stochastic Neighbor Embedding (t-SNE)^59^ algorithm built into FlowJo V10. Epitopes on BM cells from *Irf4^−/−^* and wt control mice used for dimensional reduction analysis comprised B220, sIgµ, CD43, CD24, BP-1. For RNA and WES analyses, BM cells were surface labeled for B220 and sIgµ expression and B220^+^sIgµ^-^ cells were sorted using a FACS Aria III (BD Bioscience). Sorting efficiency was routinely above 95 %. To determine cell viability, AnnexinV/propidium iodide (PI) staining was performed using 5 µL AnnexinV (640905, Biolegend) per 500 µL HBSS. After 20 min of incubation at room temperature in the dark, 1 µL propidium iodide (421301, Biolegend) was added and cells were immediately measured.

### Copy number variations (CNVs) analysis

CNVs were analyzed in tumour samples 8, 10 and 14 and compared to *Irf4^−/−^* normal tail tissue. Whole DNA was extracted from 5 x 10^6^ cells per sample using the Macherey-Nagel NucleoSpin Tissue kit (REF 740952.50) according to the manufacturer’s protocol. Library preparation was performed using the Illumina Nextera DNA kit according to manufacturer’s instruction. Sequencing was performed on an Illumina-HiSeq-1500 platform in rapid-run mode at the Genomics Core Facility of Philipps-University Marburg. Fastq quality control was performed using custom scripts. Raw sequenced reads were aligned to the Ensembl Mus musculus reference (revision 79) using Bowtie2 (version 2.0.0)^60^ with standard parametrization. Analysis of CNVs was performed using the cn.mops (Copy Number estimation by a Mixture Of PoissonS) package (version 1.18.1)^61^ with the following parametrization: prior impact = 1, lower threshold -0.9, upper threshold = 0.5 minimum width = 4. Window length was set to 10000 and the algorithm was run in unpaired mode.

### BM cryosections and analysis of B progenitor vicinity to IL-7^+^ BMSCs

Mouse femora from *Irf4^−/−^* or *wt il7^eGFP^* reporter mice were explanted, cleaned from soft tissues and fixated over-night in 4 % PFA PBS (Alfa Aesar). Samples were then dehydrated by incubation in 30 % sucrose in PBS for 24 h. Dried and dehydrated femora were snap frozen in cryomolds® (Tissue-Tek) using O.C.T freezing medium (Tissue-Tek) by being placed in a beaker of Hexan, surrounded by a beaker of Acetone and dry-ice. Samples were stored at - 20°C until processing. Cryosections of 7 µm were generated with a Leica cryostat (DB80 LX microtome blades, Leica) using Kawamoto tape^62^ (Section-lab) as described before.^63^ Cryosections were stained with antibodies against B220 (RA3-6B2, Biolegend), CD2 (14- 0021-85, eBioscience, conjugated to AF555 using lightning-Link kit, abcam), GFP (Rockland goat polyclonal anti-GFP, 600-101-215) with secondary rabbit anti-goat F(ab’)2 AF488 (thermo-scientific A21222). Samples were then mounted in DAPI ProLong Gold Antifade (ThermoFisher Scientific). Images were recorded using a Leica confocal (SP8i) microscope. Image analysis was performed in IMARIS (version 9.7.2).

### Histological analyses

Tissue samples were immediately fixed in 4 % PFA PBS solution. Histological analysis was performed on 3 µm thick sections from paraffin embedded tissue as described previously.^64^ Briefly, rehydrated paraffin sections were first blocked with 0.3 % H_2_O_2_ and goat normal serum. For immunohistochemical (IHC) stainings, rat antibodies against CD45R/B220 (clone RA3-6B2, BD) and KI67 (clone TEC-3, Dako) were then incubated on the tissue slices and bound antibody was detected with biotinylated goat anti-rat IgG (Southern Biotechnology). Bound antibody was visualized with the Vectastain-kit (Vector Laboratories) according to the manufacturer’s protocol. Hematoxylin-Eosin (HE) stainings were performed according to standard procedures. Cells of the granulocytic lineage were stained on paraffin embedded tissues with the Napthol AS-D Chloracetate (Specific Esterase, CAE) Kit (Ref: 91C-1KT, Sigma-Aldrich) according to the manufacturers protocol.

In the *in vivo* therapeutic experiments, we calculated the narrowing of the spinal cord using the equation A_t_/(A_sca_-A_sp_), where A_sca_ is the area of the spinal canal, A_t_ that of the tumor and A_sp_ that of the spinal cord area. Two different cross sections per animal were examined. The infiltration of the liver was calculated by dividing the tumor area in the liver by the whole area of the liver section. Three whole liver sections were analyzed per animal. All measurements were performed using Fijii.^65^

### Whole Exome Sequencing and biostatistical analysis

To determine single nucleotide variants (SNV) within leukemia samples, genomic (g)DNA was extracted both from primary *Irf4^−/−^* tumours as well as FACS-sorted control B220^+^mIgM^-^ BM fr.A-D cells using the High Pure PCR Template Preparation kit from Roche (11796828001). Integrity of resultant gDNA was confirmed in a 2 % Agarose gel. Macrogen in Seoul performed SureSelect All Exon V6 library preparation and sequenced exons on a NovaSeq platform producing 2 x 150 bp reads at a coverage of 100x (50x on-target coverage). Fastq quality control was performed using FASTQC (version 0.11.9). Raw sequenced reads were aligned to the Ensembl Mus musculus reference (revision 96) using STAR (version 2.6.1d) using default parametrization. Soft-clipped aligned reads were then subjected to variant calling analysis. Position-wise pile-up files were generated using samtools (version 1.9) with the mpileup option and a pileup quality threshold of 15, both for single sample and matched variant calling. Subsequently, variant calling was performed for SNP and InDel detection using VarScan2 (version 2.3.9) on single samples with the following parametrization: sampling depth = 100000, minimum variant frequency = 0.05, minimum coverage = 8, minimum variant reads = 2, minimum average read quality = 15 and a p-value threshold was set to 0.05. Only primary alignments were considered, the strand filter was enabled, and duplicates were removed. As a comparison, matched tumour-normal variant calling was performed with VarScan as well using identical parameter setting with the somatic p-value threshold set to 0.05.

For Fig.3n raw sequenced reads were aligned to the Ensembl Mus musculus reference (revision 96) using Burrows-Wheeler Aligner (BWA version 0.7.17) using default parametrization.^66^ Prior to variant calling, aligned reads were filtered using a custom filter that excludes reads with more than 3 mismatches, more than 2 indels or a mapping quality below 20 using pysam (version 0.16.0.1). Duplicates were marked and removed using Picard (GATK version 4.1.6.0).^67^ Filtered aligned reads were then subjected to variant calling analysis. Position-wise pile-up files were generated using samtools (version 1.9) with the mpileup option and a minimal base quality threshold of 20. Subsequently, variant calling performed for SNP detection using VarScan2 (version 2.4.4) using matched tumor-normal (somatic) mode with the following parametrization: sampling depth = 100000, minimum variant frequency = 0.2, minimum coverage = 8, minimum variant supporting reads = 5, minimum average read quality = 20 and a somatic p-value threshold was set to 0.05. Only primary alignments were considered, the strand filter was enabled. SNP calls were filtered to high confidence somatic mutations using VarScan’s somaticFilter method, SNPs with a variant allel frequency above 0 in the matched reference sample were excluded.

### Sanger Sequencing and polymerase chain reaction

SNVs in the JAK3 gene were confirmed by Sanger sequencing of PCR fragments spanning the *Jak3* pseudokinase and kinase region (primers used for PCR amplification and Sanger Sequencing: mJAK3 for, mJAK3 rev s. supplemental data). Sequencing services were provided by Microsynth Seqlab. To determine clonality of tumour cells, the VµH region was amplified by PCR. Amplicons were run on an agarose gel and extracted using the QIAquick Gel Extraction Kit (Qiagen). DNA fragments were then cloned into the vector pJet1.2 (Thermo Scientific) and transformed into DH10B E. coli. The indicated numbers of clones (Fig. 1G) for each PCR amplicon were sequenced and aligned with software from IMGT/V-quest.^68^

### Retroviral transduction of *Jak3-*mutants and IL-7 independency assay

The coding sequence of murine *Jak3* was amplified from pCineo-Jak3 (a gift from Olli Silvennoinen from Tampere-university in Finland) and cloned into the pMSCV-Thy1.1 expression plasmid using *BamHI* and *SalI* resctriction digestion. Site directed mutagenesis was performed following the manufacturer’s protocol using the Quick-Change II site-directed mutagenesis kit (Agilent Technologies; primers employed are listed in the supplemental materials). Viral supernatant from mutated pMSCV-Thy1.1-Jak3 constructs was produced as described previously.^58^ For viral transduction, 5 x 10^5^ IL-7 dependent primary *Irf4^−/−^* preB-I cell cultures were resuspended in 400 µL RPMI medium (D5030, Sigma-Aldrich) with 600 µL viral supernatant and 1.5 µL polybrene and spun in culture plates at 2700 RPM for 90 min at 37 °C. Cells were then replenished with conditioned medium and rested for 24 h. Transduction efficiency was measured by flow cytometry using surface staining for Thy1.1 (OX-70, Biolegend). For the IL-7 independency assay (Fig. 3 B), transduced cells were split and cultured with either recombinant murine (rm)IL-7 or 10 µg/ml neutralizing anti-IL-7 antibody (BE0048, Bio X Cell). Same protocol applied for STAT5_ca_-RV transduction. The STAT5_ca_- conctruct had been generated by EcoRI excision of STAT5A1*6 from pMSCV-STAT5A1*6-NGFR^69^ and insertion into the pMIG expression plasmid.

### RNA sequencing and biostatistical analysis

RNA extraction from primary tumour samples and FACS sorted B220^+^ mIgM^-^ pro/preB cells was performed using Trizol extraction. Quality control was performed using the Bioanalyzer RNA 6000 NanoChip (Agilent Technologies). Library preparation was performed at the Institute for Immunology, University Medical Center of the Johannes Gutenberg-University Mainz using the NEBNext Ultra Library Prep kit (New England Biolabs). For deep sequencing, the Illumina-HiSeq-4000 platform was used (Beijing Genomic Institute). Quality control on the sequencing data was performed with the FastQC tool (version 0.11.2, https://www.bioinformatics.babraham.ac.uk/projects/fastqc/). RNA sequencing reads were aligned to the ENSEMBL Mus_musculus.GRCm38 reference genome. The corresponding annotation (ENSEMBL v76) was also retrieved from ENSEMBL FTP website. The STAR aligner (version 2.4.0j) was used to perform mapping to the reference genome. Alignments were processed with the featureCounts function^70^ of the Rsubread package, using the annotation file also used for supporting the alignment. Exploratory Data Analysis was performed with the pcaExplorer package.^71^ Differential expression analysis was performed with DESeq2 package,^72^ setting the false discovery rate (FDR) cutoff to 0.1. DESeq2 datasets were analyzed using the GeneTonic^73^ and pcaExplorer packages. To assess the possible occurrence of gene fusions, we applied two different methods, Star-Fusion (version 1.10.1) and Arriba (version 2.1.0). For STAR Fusion, required meta reference files were created from the Ensembl Mus musculus reverence (revision 100) as recommended in the STAR Fusion manual. In case of Arriba, we used the mm10+GENCODEM25 assembly. In each case, we used the dockerized versions of the tools. Raw fastq files were used as an input for both tools. Subsequently, raw reads were mapped using the recommended alternative STAR settings recommended in the tools manual to leverage chimeric reads from the alignments. Default filters as recommended by the STAR-Fusion and Arriba manuals were applied to limit the false-positive rate. For the same reason, known blacklisted regions as provided by the Arriba release were excluded from the analysis.

### BCP-ALL subtype predictions using random forest classifier

Human genes (GRCh38.p13, v104) with annotated orthologous genes in mouse were extracted from ensembl database using the BiomaRt online tool. Gene counts from RNA-sequencing of a previously published human BCP-ALL cohort^34^ and of murine tumor samples were subsetted to include only human-mouse orthologous genes. Resulting gene counts were normalized by variant stabilisation transformation using the R package DESeq2 version 1.32.0. Allocation of the murine tumor samples to human BCP-ALL molecular subtypes were performed based on gene expression using a random forest machine learning algorithm (R package caret version 6.0-88) trained on the human cohort. Predictions were plotted using R package pheatmap version 1.0.12. Differential gene expression was analyzed in R package DESeq2 and resulting gene lists ranked by log2-fold-change were analyzed in GSEA version 4.1.0.

### JAK structure and sequence analysis

Mouse JAK2 (AF-Q62120) and JAK3 (AF-Q62137) structure predictions were acquired from the AlphaFold protein structure database^74^ and visualized in UCSF ChimeraX (version 1.2.5).^75^ Multiple sequence alignments were performed using the EMBL-EBI Clustal Omega tool.

### Quantitative real time (qRT-)PCR

Total RNA was extracted both from primary *Irf4^−/−^* tumors as well as FACS-sorted control B220^+^sIgµ^-^ BM fr.A-D cells of either *Irf4^−/−^* or wt animals using the Gdansk extractme kit (EM09.1) according to the manufacturers protocol. cDNA was prepared from whole RNA samples using the RevertAid cDNA kit from Thermo Fisher (K1621). qRT-PCR for *Aicda*, *Spi1* and *Pax5* was performed using the SybrGreen MasterMix reagent (4385612, AppliedBiosystems) in a StepOnePlus cycler (AppliedBiosystems). Data presented as percentage of HPRT using the formula x = 1 / 2^(cyclesAicda - cyclesHPRT)^ * 100.

### *In vivo* therapeutic studies and ultrasound imaging

Mice were injected with 3 x 10^5^ T8.1 cells intraperitoneally and monitored daily for clinical symptoms. When mice began showing signs of general morbidity, leukemia was confirmed by FACS analysis of tail vein blood for B220^+^ mIgM^-^ blast cells. When blast cells in pB reached 25 (mean 50) %, therapy was initiated with oral Dexamethasone (Jenapharm) at 6 mg/L supplied *ad libitum* in the drinking water for seven days. Maintenance therapy comprised either Ruxolitinib phosphate (S5243, Sellekchem) 1 mg (in 2% DMSO, 30% PEG300 in H_2_O, as proposed by the manufacturer), Defactinib (S7654, Sellekchem) 1.2 mg (in 5 % DMSO, 50 % PEG300, 5 % Tween 80 in H_2_O, as proposed by the manufacturer) or vehicle control (5 % DMSO, 50 % PEG300, 5 % Tween 80 in H_2_O) administered twice daily via oral gavage. During the course of disease, this treatment led to paraparesis of the hind legs and tail. A clinical scoring system was established according to the extent of paraparesis and mice were scored daily accordingly: Scores 0-3: 0) no paraparesis, 1) paraparesis induced by treatment intervention, resolves within 30 s, 2) paraparesis induced by treatment intervention, does not resolve within 30 s, 3) persistent paraparesis, independent of treatment intervention. Score 3 prompted sacrification of affected mice. High-resolution ultrasound imaging was performed using a Visual Sonics Vevo 2100 System (FUJIFILM VisualSonics, Toronto, Canada) with microscan transducer MS-550-D, 22-55MHz (FUJIFILM VisualSonics, Toronto, Canada) as described previously.^76^

### Data availability

The RNAseq datasets generated during the current study have been deposited in the Gene Expression Omnibus (GEO) archive and are available under the accession number GSE192424. The WES datasets can be accessed under PRJNA706650 in the sequencing read archive (SRA).

### Statistical analysis

Statistical analysis was performed using the GraphPad 9.0 software. Data are commonly presented as mean ± SD. Prior to significance testing, normal distribution and homogeneity of variances was confirmed by Shapiro–Wilk test and Brown–Forsythe testing. Statistical significance when comparing two normally distributed groups was evaluated using two-tailed unpaired t-tests. In case of significant differences in variances between groups, Welch’s correction was applied to account for non-norminal distribution of data. When comparing multiple groups, one-way or two-way analysis of variance (ANOVA) was performed, depending on the number of variables that differed between compared groups. This was followed by a Tukey’s Sidak, or Dunnett’s *post hoc* test, as indicated in figure legends. An alpha level of P < 0.05 was employed for significance testing.

### Study approval

All animal experiments were approved by the local government (Regierungspräsidium Gießen, G49/2018, G34/2021) and conducted according to the German animal protection law.

## Primers

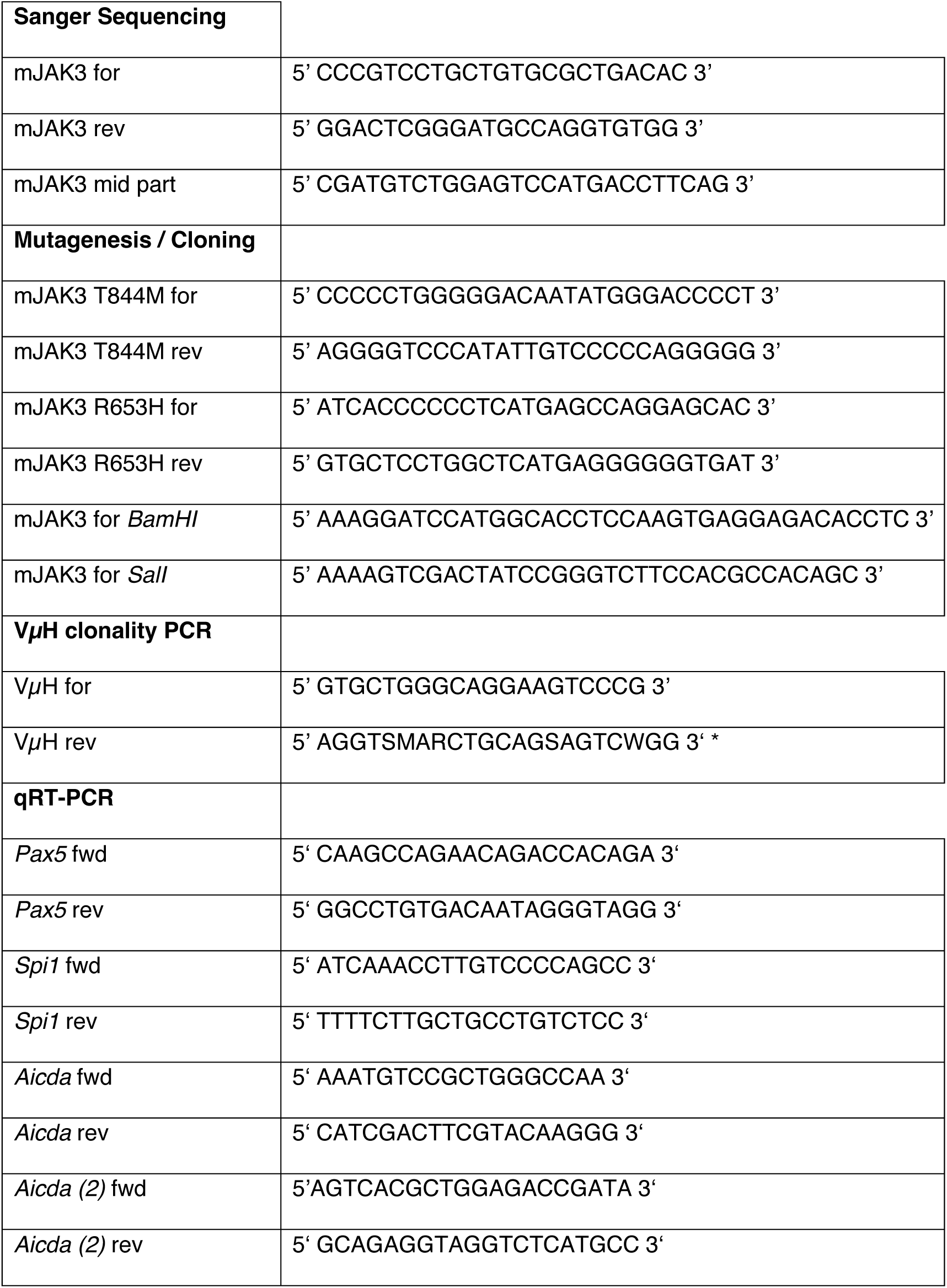

**\*:** VµH rev represents a mixture of different primers according to a degenerate nucleotide code, for amplicifation of more VµH sequences. S: C/G, M: A/C, R: A/G, W: A/T

## Author contributions

D.D.G., C.P., N.S., M.B., F.H., M.L. designed experiments; D.D.G., C.P., N.S., M.B., D.S., L.M., B.C., F.H. performed experiments; D.D.G., C.P., M.B., H.R., E.R., P.D., M.W., M.M., A.N., U.M.B., F.M., F.H., M.B., H.M.J., A.N., A.B., M.K., T.B., T.S., A.P. M.L. analyzed data; D.D.G. and M.L. prepared the manuscript.

## Acknowledgements

The authors want to thank Koji Tokoyoda (DRFZ, Berlin) for supplying *Il7^eGFP^* mice for breeding, Olli Silvennoinen (Tampere-university, Finland) for supplying us with the JAK3 construct and to Fritz Melchers (DRFZ Berlin) for the JIL-7.6 J558 and ST2 cells. Further, Hyun-Dong Chang and Anja Hauser (both DRFZ, Berlin) for supplying OP-9 cells and Kawamoto materials respectively. D.D.G. received personal funding through the German Cancer Aid, Mildred-Scheel doctoral scholarship (70112922). M.L. was funded by the Deutsche Forschungsgemeinschaft (DFG) (LO 396/8-1) and the Else Kröner-Fresenius-Stiftung.

