## Supplementary material for "IRF4 deficiency vulnerates B cell progeny for leukemogenesis via somatically acquired *Jak3* mutations conferring IL-7 hypersensitivity": suppl. information

Running title: IRF4 deficiency vulnerates preB cells for leukemogenesis

Dennis Das Gupta[1], Christoph Paul[2], Nadine Samel[1,3], Maria Bieringer[1], Daniel Staudenraus[1], Federico Marini[4], Hartmann Raifer[1], Lisa Menke[1], Lea Hansal[1], Bärbel Camara[1], Edith Roth[5], Patrick Daum[5], Michael Wanzel[6], Marco Mernberger[6,7], Andrea Nist[6], Uta-Maria Bauer[6], Frederik Helmprobst[8,9], Malte Buchholz[10], Katrin Roth[11], Lorenz Bastian[12], Alina M Hartmann[12], Claudia Baldus[12], Koichi Ikuta[13], Andreas Neubauer[14], Andreas Burchert[14], Hans-Martin Jäck[5], Matthias Klein[15], Tobias Bopp[15,16], Thorsten Stiewe[6,7], Axel Pagenstecher[8,9], Michael Lohoff[1]

[1] *Institute for med. Microbiology & Hospital Hygiene, Philipps University Marburg, Germany*

[2] *University Hospital Gießen and Marburg, and Philipps University, Dept. Ophthalmology, Marburg, Germany*

3MVZ for Laboratory Medicine and Microbiology, Koblenz-Mittelrhein, Germany

[4] Institute of Medical Biostatistics, Epidemiology and Informatics (IMBEI), University Medical Center of the Johannes Gutenberg-University Mainz, Germany

[5] *Division of Molecular Immunology, Nikolaus-Fiebiger Center, University of Erlangen-Nürnberg, Erlangen, Germany*

[6] Institute for Molecular Biology and Tumor Research (IMT), Center for Tumor- and Immunobiology (ZTI), Philipps University Marburg, Germany

[7] Genomics Core Facility, Philipps University Marburg, Germany

[8] Core Facility for Mouse Pathology and Electron Microscopy, Philipps University Marburg, Germany

[9] University Hospital Gießen and Marburg, *and Philipps University,* Institute of Neuropathology, Marburg, Germany

[10] *University Hospital Gießen and Marburg*, and Philipps University, Clinic for Gastroenterology and Core Facility Small Animal Ultrasound, Marburg, Germany

[11] Core facility for Cellular Imaging, Philipps University Marburg, Germany

[12] Medical Department II, Hematology and Oncology, University Medical Center Schleswig- Holstein, Kiel, Germany

[13] Institute for Frontier Life and Medical Sciences, Kyoto University, Japan

[14] *University Hospital Gießen and Marburg, and Philipps University, Dept. Hematology, Oncology and Immunology, Marburg,* Germany

[15] Institute for Immunology, Research Center for Immunotherapy (FZI), University Cancer Center, University Medical Center of the Johannes Gutenberg-University Mainz, Germany

[16] German Cancer Consortium (DKTK)

The authors have declared that no conflict of interest exists.

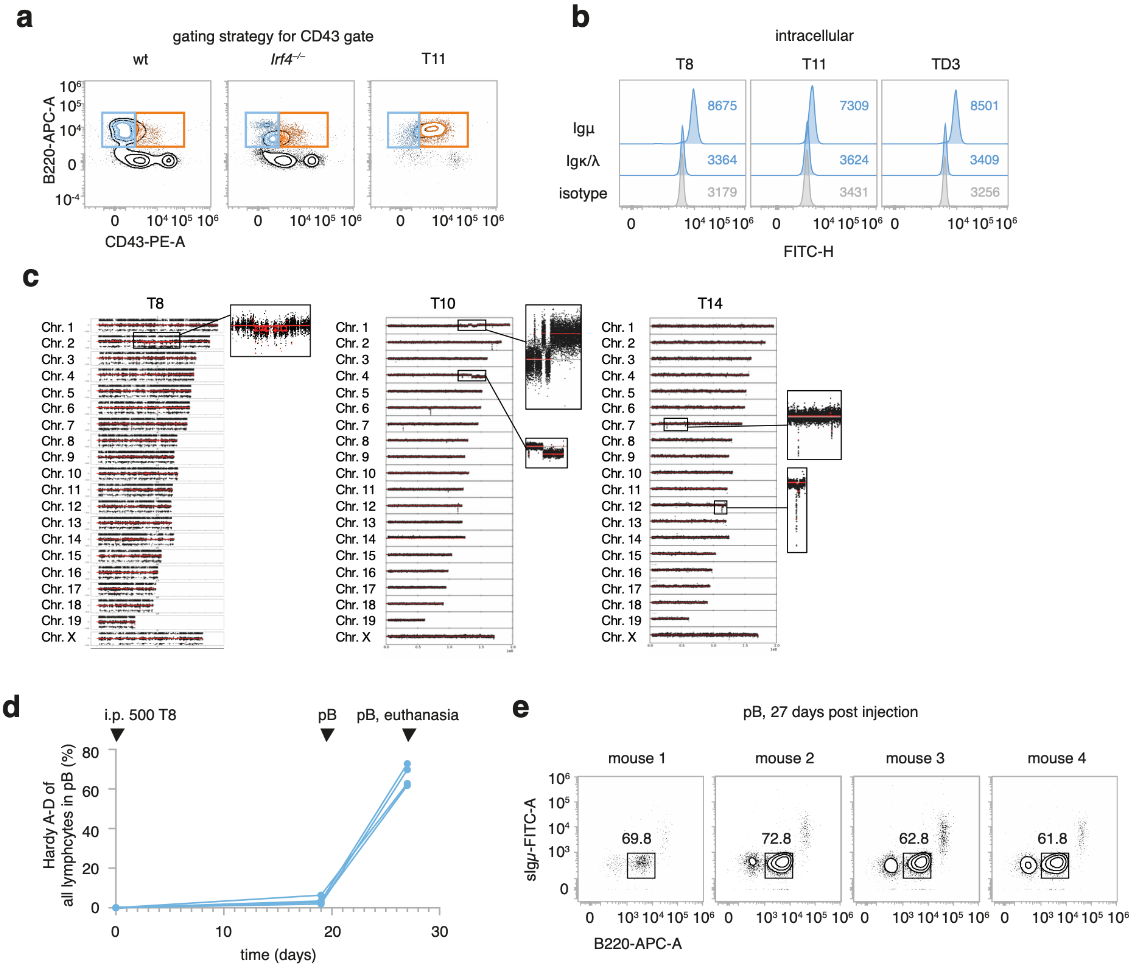


**Supplementary Fig.1: extended data to Fig.1**

**a** gating strategy for Fig.1j: wt and *Irf4*$^{-/-}$ control BM stainings were used to identify the CD43$^{+}$ cell gate used for tumour cell phenotyping **b** T8, T11 and TD3 were stained for intracellular Igµ and light chain (Igκ/λ) expression or isotype control. Numbers indicate geometric mean fluorescence intensity. **c** gDNA (T10, T14) or exome (T8) libraries were sequenced and analyzed using the cn.mops pipeline for copy number variations (CNV); inserts highlight altered chromosomal regions. **d** four wt mice were *i.p*. injected with 500 T8 cells. After 19- and 27-days tail vein blood (pB) was analyzed by flow cytometry for the presence of B220$^{mid}$ Igµ$^{-}$ leukemia cells. Dots represent individual mice. **e** flow cytometric analysis of pB for of B220 and sIgµ at 27 days post injection. Numbers indicate frequency within the respective gate (%).

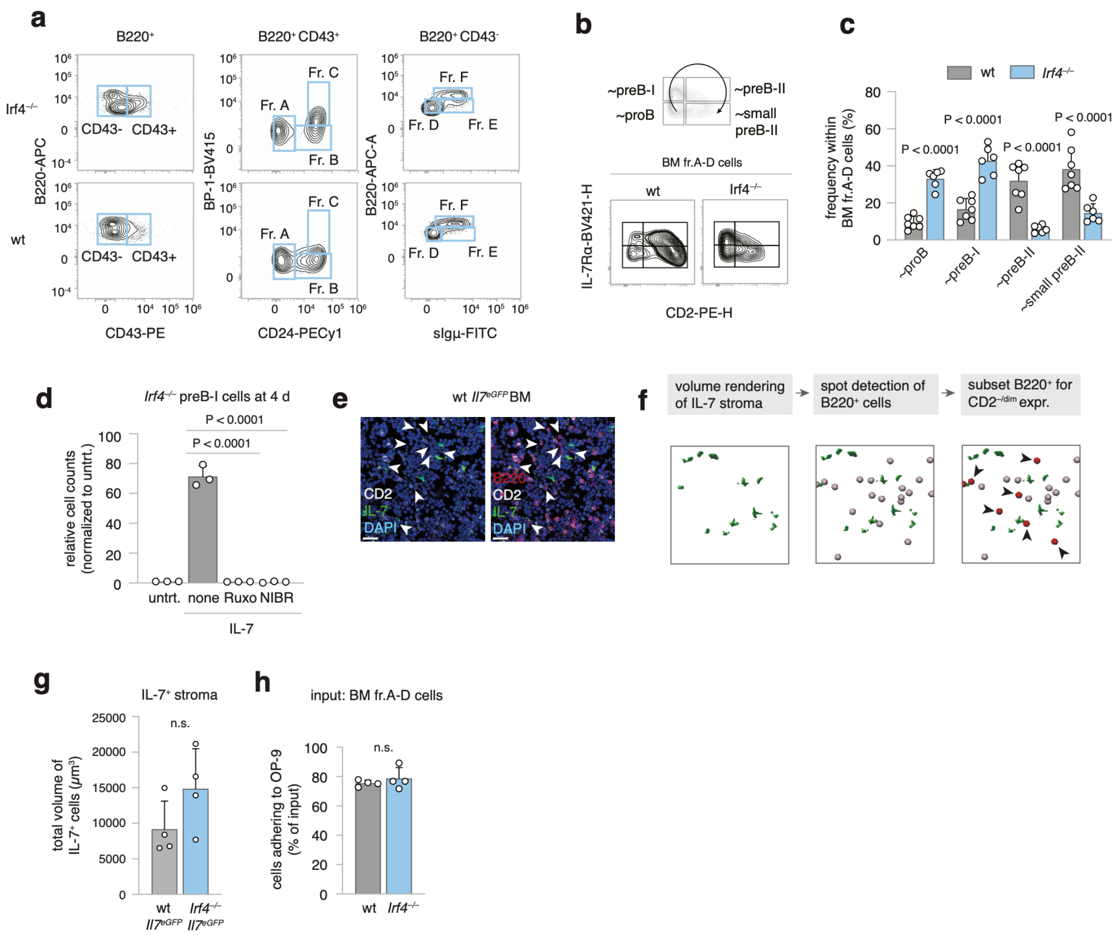


**Supplementary Fig.2: extended data to Fig.2**

**a** Gating strategy for Hardy fragment analysis in Fig.2a-f. Gated B220$^+$, B220$^+$CD43$^+$ and B220$^+$CD43$^-$ were analyzed for expression of the indicated markers. **b** within the fr.A-D cell gate, BM cells were analyzed for CD2 and IL-7Rα expression. Top pictogram presenting developmental path within the gated quadrants. **c** Quantification of cell frequency within the fr.A-D cell gate (**b**) for n = 7 (wt) and n = 6 (*Irf4*$^{-/-}$) mice. Analyzed with two-way ANOVA, Sidak post-hoc **d** *Irf4*$^{-/-}$ BM cells were cultured for 4 days in the presence of the indicated substances and cell counts recorded, presented as normalized to untrt. = untreated. Ruxo = Ruxolitinib, NIBR = NIBR3049. One-way ANOVA, Tukey post-hoc **e** Overlay confocal microscopic images of wt *il7*$^{eGFP}$ femur cryosections. Arrowheads indicate B220$^+$CD2$^{-/dim}$ cells. Scale bars = 20 $\mu$m **f** strategy for comparing B cell progenitor proximity to IL-7$^+$ BMSCs in BM cryosections using the IMARIS software (see methods). Briefly, GFP$^+$ signal was rendered as volumes, B220$^{+}$ cells were detected as spots and subsetted (red) for CD2$^{-/dim}$ expression. Distances were measured from each spot to the nearest GFP$^{+}$ volume surface. Arrowheads = B220$^{+}$, CD2$^{-/dim}$ cells. See also supplementary movie 1. **g** total volume of GFP$^{+}$ cellular structures was measured for each cryosection (n = 4 per genotype). Each dot represents one cryosection from one biological replicate. Unpaired two-tailed t-test. **h** fr.A-D cells were MACS purified from BM and seeded onto monolayers of OP-9 cells. Non-adhering cells were counted after 1 hours. Adherent cell counts calculated as $n_{adherent} = n_{input} - n_{non-adherent}$. Data depicts $n_{adherent}$ / $n_{input}$ as percentages. Unpaired two-tailed t-test. Bars represent mean ± SD in all panels.

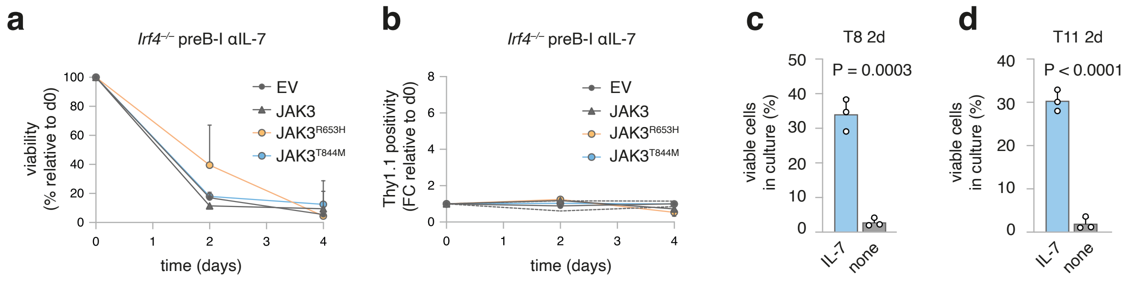


**Supplementary Fig.3: extended data to Fig.3**

**a** *Irf4*$^{-/-}$ preB-I cell cultures from BM cells were transduced with RV as in Fig.2e-g and seeded in the presence of anti(α)-IL-7. Viability of cells is plotted over the course of 4 days relative to viability at day 0. **b** Thy1.1$^{+}$ cell frequency relative to day 0 is plotted for cells as in (**a**). Data as mean ± SD of n = 3 independent experiments for (**a-b**). **c** T8 and **d** T11 cells were cultured for 48 h in the presence or absence of 10 ng/mL IL-7 and viability of culture was recorded. Dots indicate n = 3 independent experiments summarized as bars (mean ± SD), two-tailed unpaired t-test.

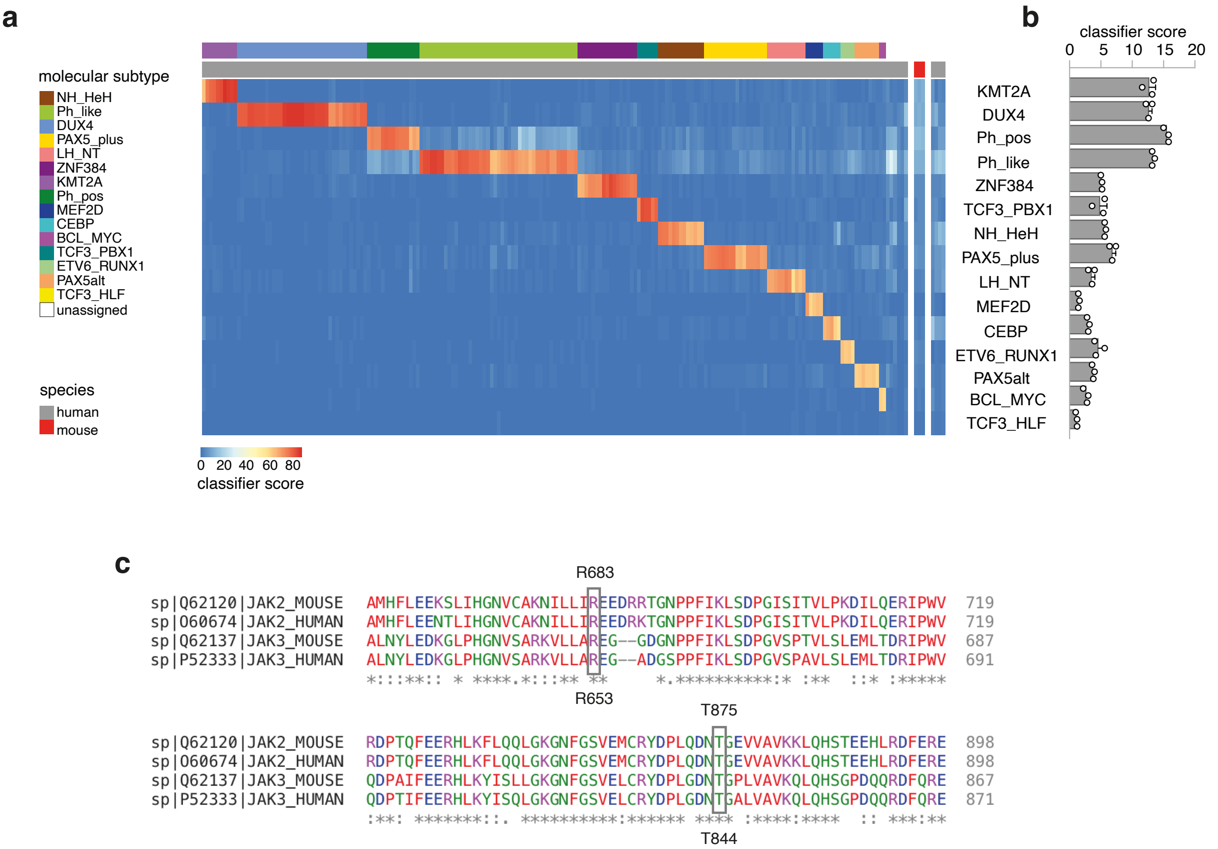


**Supplementary Fig.4: extended data to Fig.4**

**a** expression of human-mouse orthologue genes was compared between a published BCP-ALL patient cohort and *Irf4–/–* leukemia samples. A random forest classifier trained on the human samples was used to generate BCP-ALL subtype prediction scores for *Irf4–/–* leukemia samples. Heatmap presents individual patient/mouse samples as columns, ALL subtypes as rows, colour-fills give prediction scores per row. Colour labels above heatmap indicate the molecular BCP-ALL subtype of each human sample (first row) and the species (human/mouse, second row) as explained to the left of the heatmap. *Irf4–/–* leukemia samples are separated from human samples by white space. **b** classifier scores for *Irf4–/–* leukemia are plotted adjacent to the corresponding row in (**a**) for the n = 3 tested tumours, indicated by dots, summarized by bars (mean ± SD) **c-d** Clustal omega sequence alignment of mouse and human JAK2 and JAK3 **c** JH2 and **d** JH1 loop regions in mouse and human. R683/R653 and T875/T844 amino acids are highlighted.

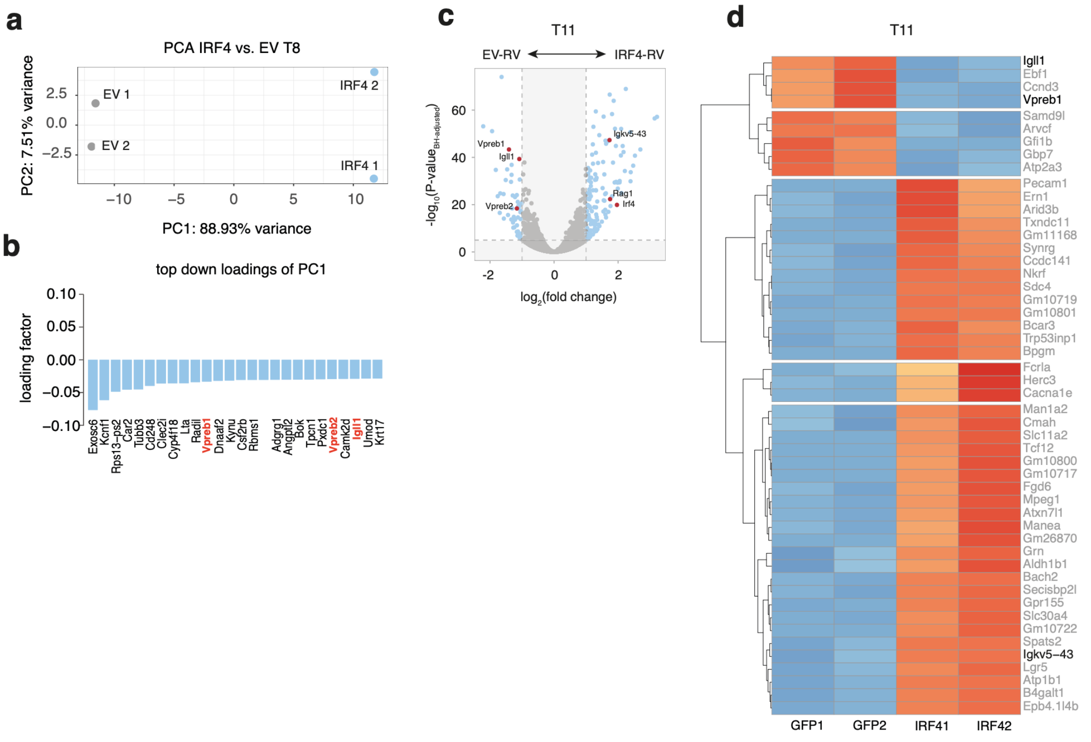


**Supplementary Fig.5: extended data to Fig.5**

**a** Principal components (PC) 1 and 2 are presented for expression data from Fig.5d-h. **b** top down-pointing loadings for PC1 are presented and ψL components highlighted. **c-d** T11 cells transduced with RVs as described in Fig.5. Total RNA sequenced 24 h after transduction. **c** Volcano plot: log$_2$ of fold change between conditions (IRF4-RV vs. EV-RV) against respective -log$_{10}$ of Benjamini-Hochberg adjusted p-values. ψL components and the differentiation genes *Igkv*, *Rag1* and *Irf4* are highlighted. Dotted lines mark x- and y- cutoff values. **E** Heatmap depicting the 50 most significantly regulated genes. Euclidean clustering depicted as dendrograms to the left. Two samples per group.

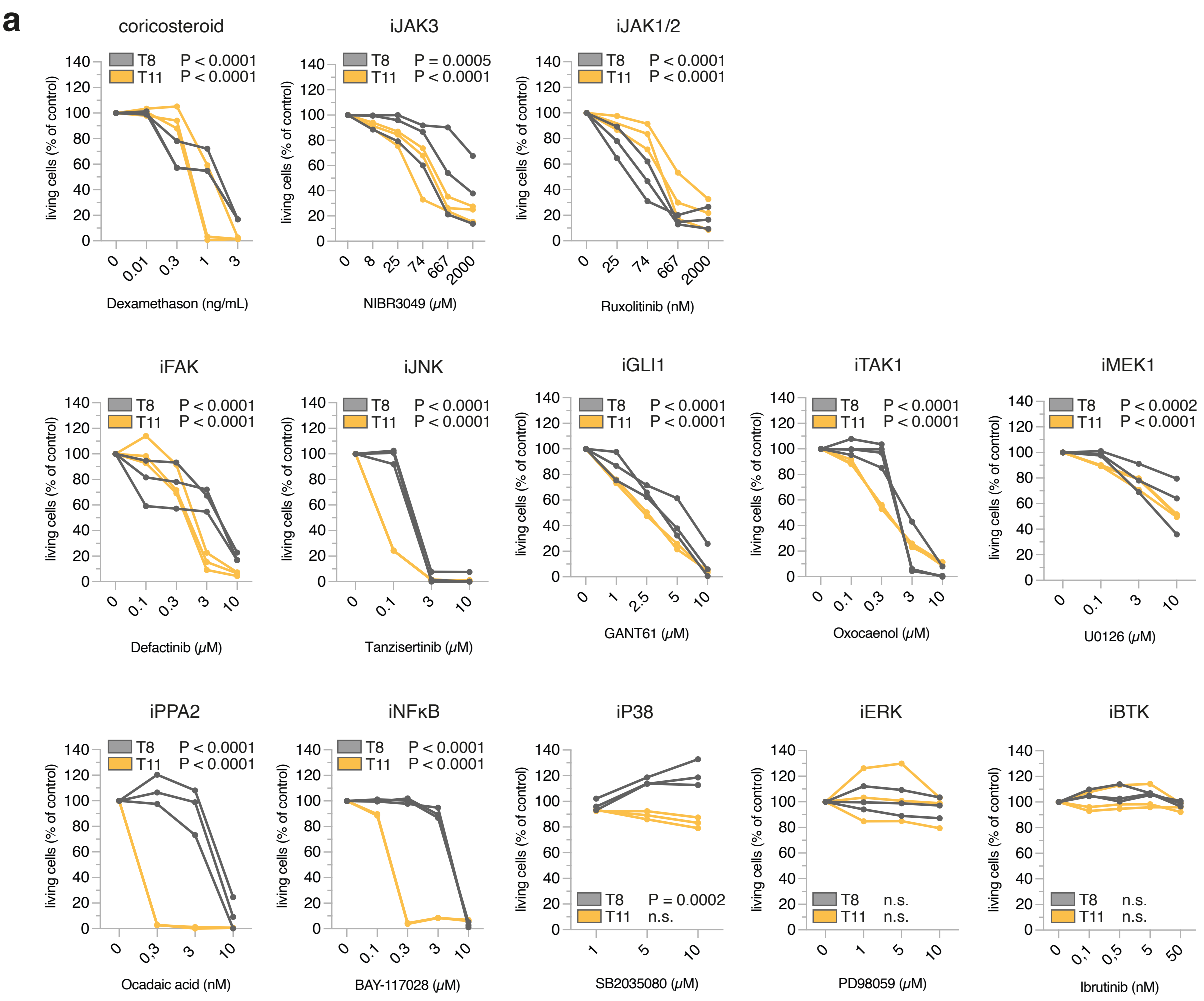


**Supplementary Fig.6: small compound inhibitors targeting leukemia cell survival**

**a** 2.5x10$^5$ T8.1 or T11 cells per well were cultured in a 48 well plate in the presence of the indicated concentrations of inhibitors listed below the x-axis. The inhibitors target the pathways described above of the panels (exception: dexamethasone). To determine the percentage of viable cells, samples were stained using Annexin V and propidium/iodide after 48 h. All values are relative to initial viability at the onset of the experiment which was arbitrarily set to 100 %. Results give the mean ± SD of at least three separate experiments per inhibitor. Unpaired two-tailed t-tests comparing viability at highest to lowest concentration of inihibitor.

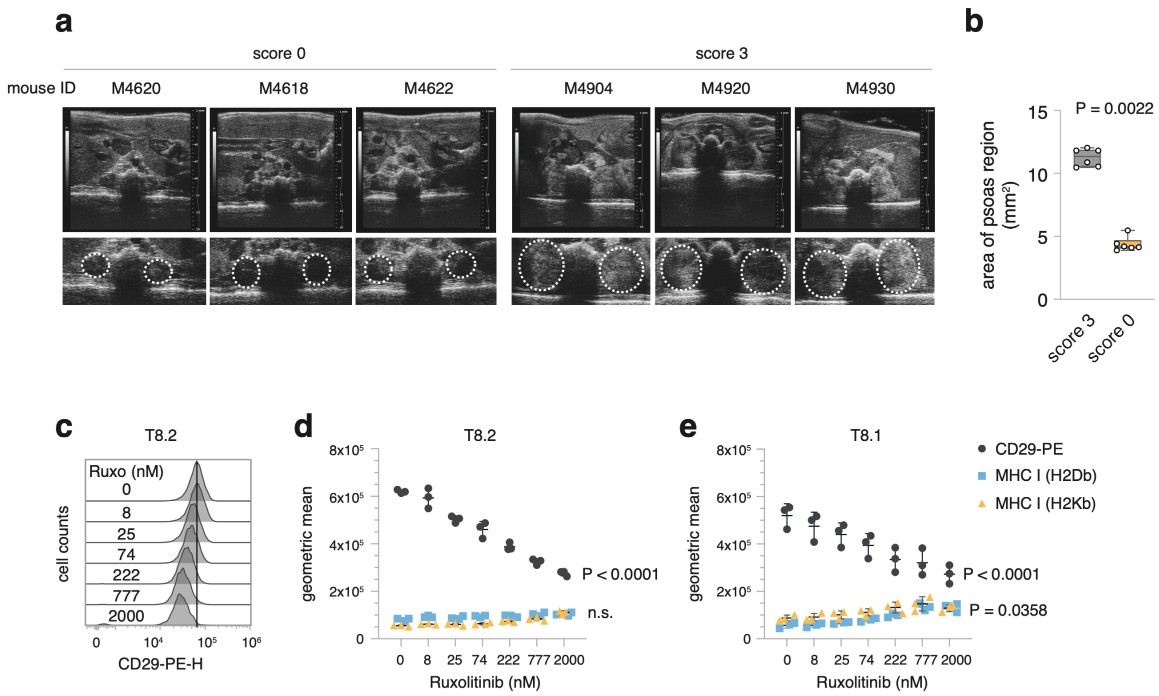


**Supplementary Fig.7: extended data to Fig.6**

**a** Ultrasound scan of the paravertebral lumbar area of individual (see mouse ID) tumour-affected mice. The region of the psoas muscle is highlighted as a circle, the area of which is measured. **b** Bars show the mean area ± SD of the psoas region for three mice with score 0 and 3, analyzed by two-tailed unpaired t-test. **c-e** T8.1 and T8.2 cells were cultured for 48 h in the presence of varying concentrations of Ruxolitinib. Thereafter, CD29, MHC I (H2Db and H2Kb) surface expression were measured. **c** Representative histogram for CD29 expression on T8.2 exposed to Ruxolitinib. **d** Quantification of results for T8.2, **e** Quantification for T8.1. Two-Way ANOVA, Sidak post-hoc comparisons to data for 0 nM Ruxolitinib.

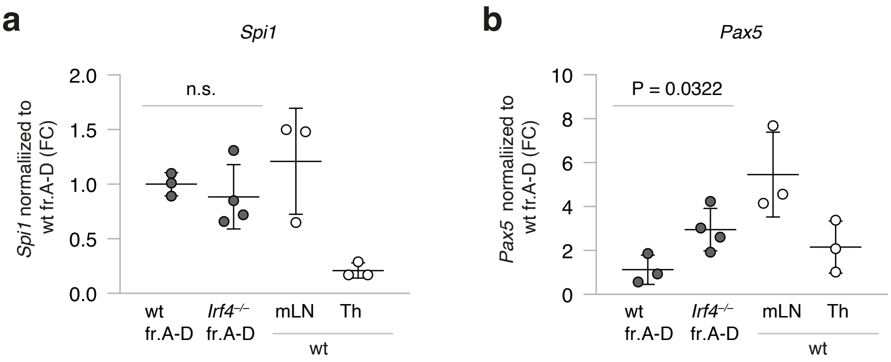


**supplementary Fig.8: *Pax5* and *Spi1* are not downregulated in *Irf4*$^{-/-}$ fr.A-D cells**

**a** *Spi1* and **b** *Pax5* gene expression in sorted *Irf4*$^{-/-}$ and wt fr.A-D cells, as well as wt mLN and T helper (Th) control samples. n = 3 biological replicates for all but *Irf4*$^{-/-}$ fr.A-D cells (n=4). Mean ± SD, dots represent individual samples, Two-tailed unpaired t-test comparing wt to *Irf4*$^{-/-}$ fr.A-D cells

| | samples | | |
|---|---|---|---|
| | T7 | T8 | T12 |
| number of independent sequencing reactions | 27 | 30 | 28 |
| number of successful sequencing reactions | 24 | 29 | 13 |
| number of identical sequences | 23 | 28 | 12 |

**Supplementary Table 1: Sequencing of VDJ junctions for IgH reveals that tumour samples are clonal**

| | characteristics | | | lifespan | | | *Jak3* mutation | | | | |
|---|---|---|---|---|---|---|---|---|---|---|---|
| tumour | genotype | type | sex | birth (dd/mm/yy) | death (dd/mm/yy) | age (days) | SNP | method | Reference Base | Alternative Base | frequency |
| T8 | C57BL/6 $Irf4^{-/-}$ | primary | m | 09/08/10 | 06/04/11 | 240 | p.R653H | WES | G | A | 0.50 |
| T10 | C57BL/6 $Irf4^{-/-}$ | primary | m | 19/07/10 | 12/04/11 | 267 | p.T844M | WES | C | T | 0.57 |
| T11 | C57BL/6 $Irf4^{-/-}$ | primary | m | 24/10/10 | 11/05/11 | 199 | p.R653H | WES | G | A | 0.46 |
| T14 | C57BL/6 $Irf4^{-/-}$ | primary | m | 24/08/10 | 06/07/11 | 316 | p.L634W | RNA | T | G | 0.50 |
| TD1 | C57BL/6 $Irf4^{-/-}$ | primary | w | 30/09/17 | 07/06/18 | 250 | p.L627F | Sanger | G | A | 0.50 |
| TD2 | C57BL/6 $Irf4^{-/-}$ | primary | w | 30/09/17 | 08/06/18 | 249 | p.A742V | Sanger | G | A | 0.50 |

**Supplementary Table 2: *Jak3* mutations identified in five out of five tested leukemia samples**
